# Neural populations in the language network differ in the size of their temporal receptive windows

**DOI:** 10.1101/2022.12.30.522216

**Authors:** Tamar I. Regev, Colton Casto, Eghbal A. Hosseini, Markus Adamek, Anthony L. Ritaccio, Jon T. Willie, Peter Brunner, Evelina Fedorenko

## Abstract

Despite long knowing what brain areas support language comprehension, our knowledge of the neural computations that these frontal and temporal regions implement remains limited. One important unresolved question concerns functional differences among the neural populations that comprise the language network. Leveraging the high spatiotemporal resolution of intracranial recordings, we examined responses to sentences and linguistically degraded conditions and discovered three response profiles that differ in their temporal dynamics. These profiles appear to reflect different temporal receptive windows (TRWs), with average TRWs of about 1, 4, and 6 words, as estimated with a simple one-parameter model. Neural populations exhibiting these profiles are interleaved across the language network, which suggests that all language regions have direct access to distinct, multi-scale representations of linguistic input—a property that may be critical for the efficiency and robustness of language processing.

## Introduction

Language processing engages a network of brain regions that reside in the temporal and frontal lobes and are typically left-lateralized (e.g., Fedorenko et al., 2010; Pallier et al., 2011). These brain regions respond strongly to linguistic stimuli across presentation modalities (Fedorenko et al., 2010; Vagharchakian et al., 2012; Regev et al., 2013; Scott et al., 2017), tasks (Fedorenko et al., 2010; Cheung et al., 2020; Diachek, Blank, Siegelman et al., 2020), and languages (Malik-Moraleda, Ayyash et al. 2022). This language-responsive network is highly selective for language, showing little or no response to diverse non-linguistic inputs and tasks (e.g., Fedorenko et al., 2011; Monti et al., 2012; Deen et al., 2015; Ivanova et al., 2020, 2021; Liu et al., 2020; Chen et al., 2023; Shain, Paunov, Chen et al., 2023; see Fedorenko, Ivanova & Regev, in press, for a review). However, the precise computations and neuronal dynamics that underlie language comprehension remain debated.

Based on neuroimaging and aphasia evidence, some have argued for dissociations among different aspects of language, including phonological/word-form processing (e.g., Okada and Hickok, 2006; Graves et al., 2008; DeWitt and Rauschecker, 2012), the processing of word meanings (e.g., Price et al., 1997; Rodd et al., 2005; Mesulam et al., 2013), and syntactic/combinatorial processing (e.g., Friederici, 2002, 2011; Hagoort, 2005; Grodzinsky and Santi, 2008; Matchin and Hickok, 2020). However, other studies have reported distributed sensitivity to these aspects of language across the language network (Fedorenko et al., 2010, 2020; Bautista and Wilson, 2016; Blank et al., 2016; Anderson et al., 2021; Caucheteux et al., 2021; Reddy & Wehbe, 2021; Shain, Blank et al., 2020; Regev et al., 2024). Some of the challenges in discovering robust functional differences within the language network may have to do with the limitations of fMRI—the dominant methodology available for studying language processing. Each fMRI voxel contains a million or more individual neurons, which may differ functionally. If different linguistic computations are implemented in distinct neural populations that are distributed and interleaved across the language cortex, such dissociations may be difficult to detect with fMRI. Further, the relatively slow temporal resolution of fMRI (typically, ∼2 seconds) may obscure the dynamics of linguistic computations.

In recent years, invasive recordings of human neural activity (e.g., Mukamel and Fried, 2011), including electrocorticography (ECoG) and stereo electroencephalography (sEEG), have become increasingly available to language neuroscience researchers, as patients undergoing presurgical evaluation (usually for intractable epilepsy) agree to perform linguistic tasks while implanted with intracranial electrodes. These data have high spatial and temporal resolution, allowing the tracking of neural dynamics across both space and time. Several previous studies have probed intracranial neural responses during language comprehension (e.g., Fedorenko et al., 2016; Nelson et al., 2017; Woolnough et al., 2023; Desbordes et al., 2023; Goldstein et al., 2022; 2023). For example, Fedorenko et al. (2016) reported sensitivity in language-responsive electrodes to both word meanings and combinatorial processing, in line with fMRI findings (e.g., Fedorenko et al., 2010; Bedny et al., 2011). They also reported a temporal profile where neural activity gradually increases (builds up) across the sentence (replicated by Nelson et al., 2017; Desbordes et al., 2023; Woolnough et al., 2023), which they interpreted as reflecting the construction of a sentence meaning. However, considerable disagreement exists in the field regarding the number of distinct profiles that characterize cortical language responses, how they functionally differ, and what computations they collectively support in the service of language comprehension and production.

Here, we report a detailed investigation of neural responses during language processing. To isolate the language network from nearby lower-level perceptual areas and domain-general cognitive areas, we focus on electrodes that show a characteristic functional signature of the language areas: a stronger response to sentences than to sequences of nonwords (as in Fedorenko et al., 2016). To foreshadow our findings, we report three response profiles that differ in their temporal dynamics and overall magnitude of response to linguistically degraded conditions. Using a toy model with a single parameter—the timescale of information integration—we argue that these profiles reflect distinct temporal receptive window sizes in the language system (e.g., Lerner et al., 2011; Blank and Fedorenko, 2020; Jain et al., 2020).

## Results

We used intracranial recordings from patients with intractable epilepsy to investigate neural responses during language comprehension. Participants in Dataset 1 were presented with four types of linguistic stimuli that have been traditionally used to tease apart neural responses to word meanings and syntactic structure (Fedorenko et al., 2010, 2012, 2016; Pallier et al., 2011; Shain, Kean et al., in press; Desbordes et al., 2023; for earlier uses of this paradigm, see Mazoyer et al., 1993; Friederici et al., 2000; Humphries et al., 2001; Vandenberghe et al., 2002): sentences (S), lists of unconnected words (W), Jabberwocky sentences (J), and lists of unconnected nonwords (N) (**Figure 1A-B**, <u>Methods</u>, all stimuli are available at osf.io/xfbr8/). In each trial, 8 words or nonwords were presented on a screen serially and participants were asked to silently read them. To maintain alertness, after each trial, participants judged whether a probe word/nonword had appeared in that trial. See <u>Methods</u> for further details of stimulus presentation and behavioral response data. In Dataset 2, just two of these conditions were used: sentences and lists of nonwords.

**Figure 1.**
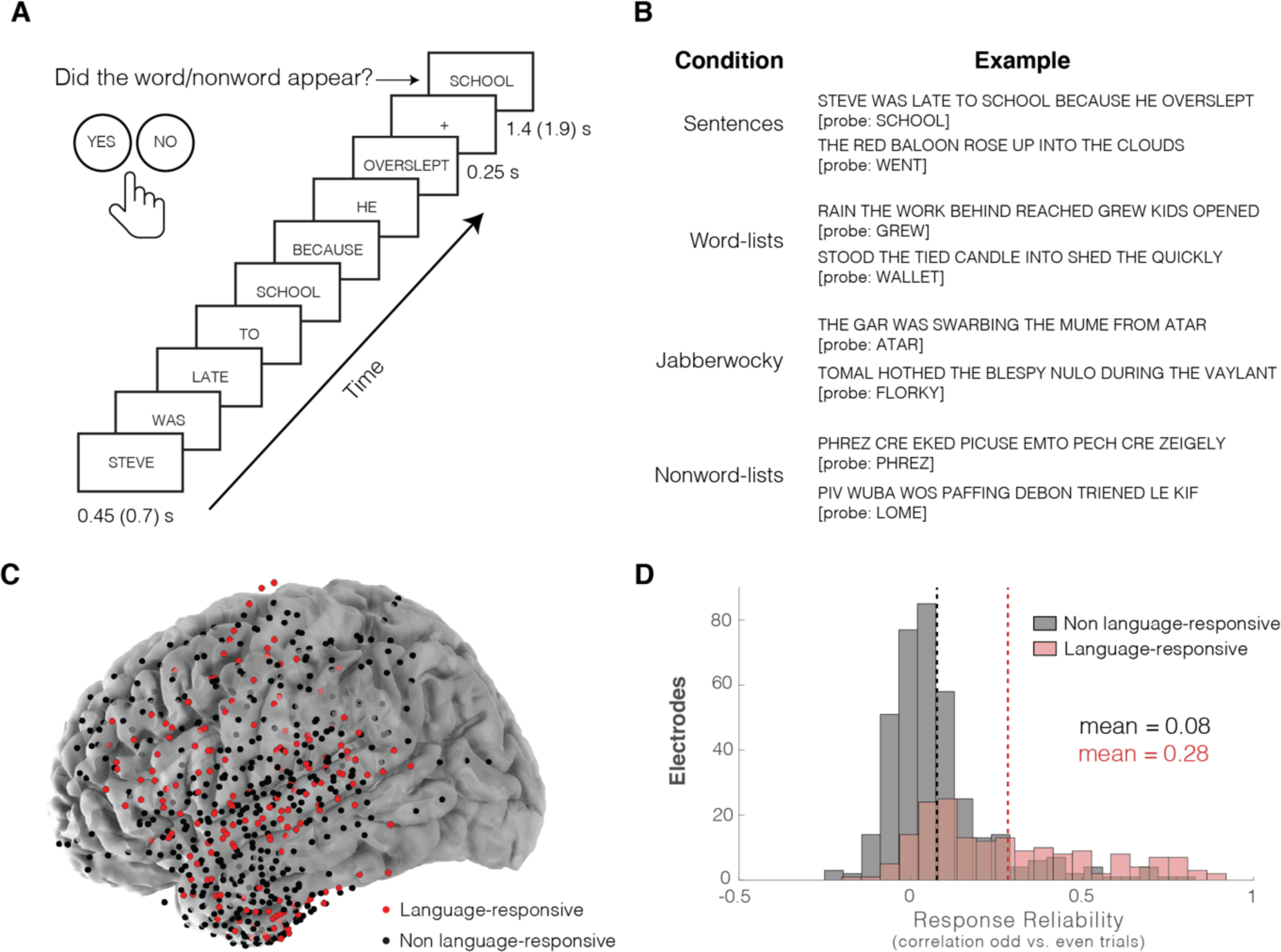
– Experimental procedure and the distribution of the implanted electrodes for Dataset 1. **A**) A sample trial from the Sentence condition. **B)** For each of the four experimental conditions, items are either presented with probes that appeared in the trial or not. Adapted from Fedorenko et al. (2016). **C)** The locations of language-responsive (n=177, red; <u>Methods</u>) and non-language-responsive (n=373, black) electrodes across the six participants in Dataset 1. Electrodes were implanted almost exclusively in the left hemisphere for Dataset 1 and concentrated in the temporal and frontal lobes. **D)** Response reliability across odd and even trials (based on a correlation of mean condition-level responses) for language-responsive and non-language-responsive electrodes. Language-responsive electrodes exhibit more reliable responses to linguistic stimuli than non-language-responsive electrodes.

We asked three research questions: 1) Does the language network contain reliably distinct response profiles? If so – 2) What do these profiles reflect? And finally – 3) Do electrodes exhibiting different response profiles tend to be located in particular regions of the language network? We used Dataset 1 (n=6) for initial evaluation of these questions because this dataset contained a richer set of experimental conditions. We then used Dataset 2 (n=16) as an attempt to replicate the findings despite the more compact experimental paradigm.

### 1. Language-responsive electrodes exhibit reliably distinct response profiles

We clustered the high gamma neural response patterns of language-responsive electrodes from Dataset 1 (6 participants, same as those used in Fedorenko et al., 2016, 177 language-responsive electrodes; **Figure 1C**, <u>Methods</u>, **Table 1**) to sentences (S), word lists (W), Jabberwocky sentences (J) and nonword lists (N) (**Figure 1A-B**). We focused on differences across experimental conditions and therefore clustering was performed on the average condition timecourses, which were concatenated across the four conditions to create a single timecourse per electrode (**Figure 2B**, <u>Methods</u>). The k-medoids clustering algorithm, combined with the “elbow” method (<u>Methods</u>), suggested that three clusters (k=3) optimally explain the data (**Figure 2A**; similar results emerged with a k-means clustering algorithm, see OSF osf.io/xfbr8/). Although we combined the electrodes from all 6 participants for clustering, electrodes that belong to each of the three clusters were evident in every participant individually (**Figure 2B**, **S1**).

**Figure 2.**
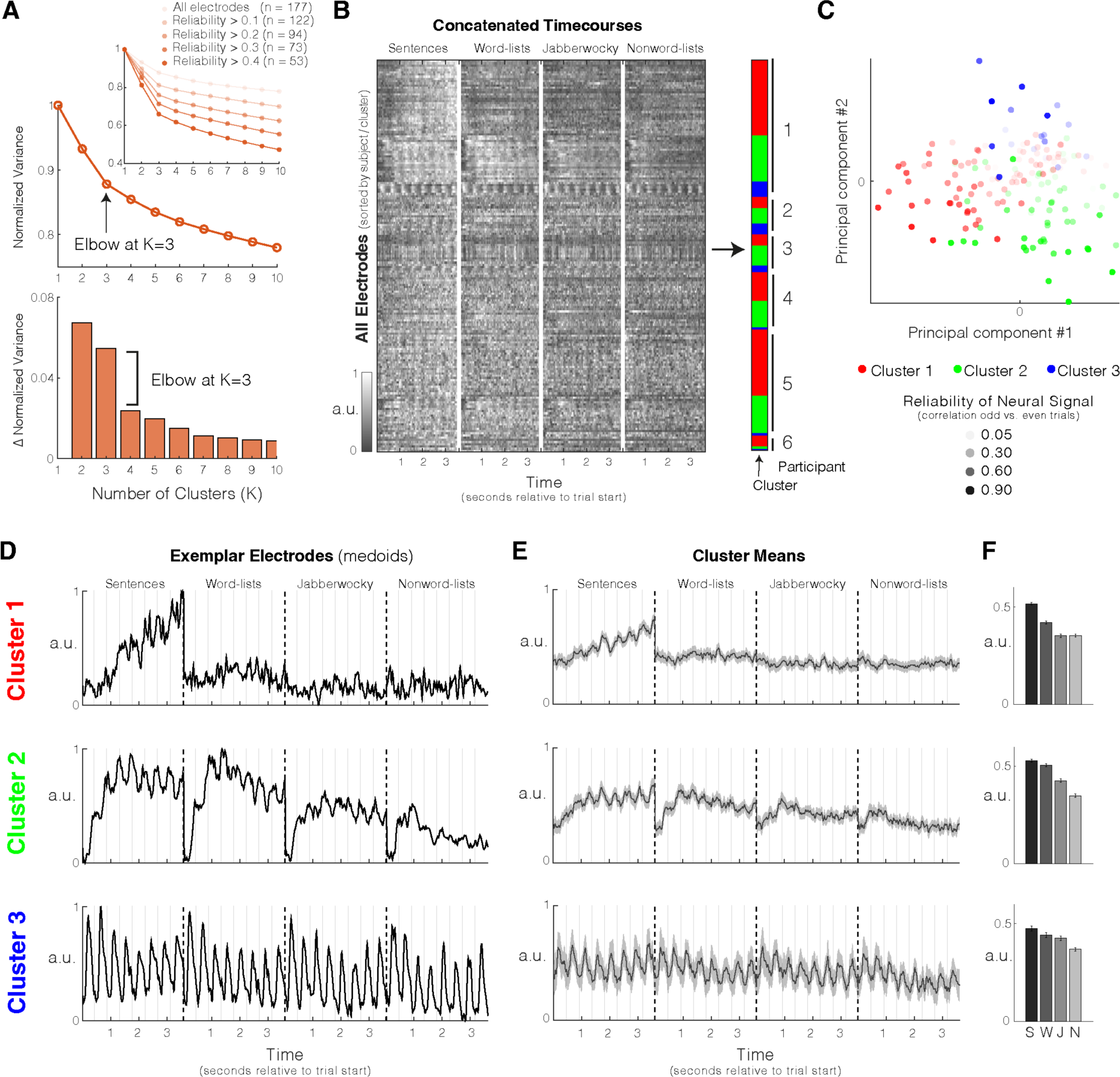
– Dataset 1, k-medoids clustering with k=3. **A**) Search for optimal k using the “elbow method”. ***Top*:** Variance (sum of the distances of all electrodes to their assigned cluster center) normalized by the variance when k=1 as a function of k (normalized variance (NV)). ***Inset***: Clustering was performed while omitting electrodes below a parametrically sampled reliability threshold. Orange shading represents the reliability threshold for omitting electrodes. The elbow (point of transition between a steeper to a more moderate slope) gets more pronounced when eliminating lower-reliability electrodes, which suggests that k=3 best describes these data. ***Bottom*:** Change in NV as a function of k (NV(k+1) – NV(k)). After k=3, there was a large drop in the change in variance. **B)** Clustering mean electrode responses (concatenated across the four experimental conditions: sentences (S), word lists (W), Jabberwocky (J), nonword lists (N)) using k-medoids (k=3) with a correlation-based distance (<u>Methods</u>). Shading of the data matrix reflects normalized high-gamma power (70-150Hz). Electrodes are sorted vertically due to participant and their assignment to clusters (right color bar). All three clusters are present in each of the six participants. **C)** Electrode responses visualized on their first two principal components, colored by cluster and shaded by the reliability of the neural signal as estimated by correlating responses to odd and even trials (**Figure 1D**). **D)** Timecourses of best representative electrodes (‘medoids’) selected by the algorithm from each of the three clusters. The timecourses reflect normalized high-gamma (70-150Hz) power averaged over all trials of a given condition. a.u. stands for arbitrary units; the signals were z-scored and normalized to have minimum value of 0 and maximum value of 1. **E)** Timecourses averaged across all electrodes in each cluster. Shaded areas around the signal reflect a 99% confidence interval over electrodes. **F)** Mean condition responses by cluster. Error bars reflect standard error of the mean over electrodes. After averaging across time, response profiles are not as distinct by cluster (especially for Clusters 2 and 3), which underscores the importance of temporal information in elucidating this grouping of electrodes.

**Table 1.** – Details for Dataset 1. (All data were collected at the Albany Medical Center (Site=AMC).) Here and in Table 2, ‘Total electrodes’ excludes reference electrodes, ground electrodes, microphone electrodes, trigger electrodes, skull EEG electrodes, and EKG electrodes; and ‘Total clean electrodes’ excludes electrodes with significant line noise, significant interictal discharges, or large visual artifacts identified through manual inspection. ‘Elec per amp’ – Number of electrodes per amplifier. ‘Pres rate (per word)’ – duration of presentation of each single word or nonword.

| Participants | Age | Sex | Site | ECoG or sEEG | Language-responsive electrodes (S>N) | Total clean electrodes | Total electrodes | Native sampling freq (Hz) | Elec per amp | Runs | Pres. rate (per word) | Trials per cond |
| --- | --- | --- | --- | --- | --- | --- | --- | --- | --- | --- | --- | --- |
| Participant 1 | 29 | F | AMc | ECoG | 62 (0 RH) | 108 (0 RH) | 120 (0 RH) | 1200 | 16 | 10 | 450ms | 80 |
| Participant 2 | 25 | F | AMc | ECoG | 17 (0 RH) | 115 (0 RH) | 128 (0 RH) | 1200 | 16 | 10 | 700ms | 60 |
| Participant 3 | 18 | F | AMc | ECoG | 17 (0 RH) | 92 (0 RH) | 98 (0 RH) | 1200 | 16 | 10 | 700ms | 60 |
| Participant 4 | 28 | M | AMc | ECoG | 26 (0 RH) | 106 (0 RH) | 134 (0 RH) | 1200 | 64 | 10 | 700ms | 60 |
| Participant 5 | 25 | F | AMc | ECoG | 48 (0 RH) | 93 (0 RH) | 98 (0 RH) | 1200 | 64 | 10 | 450ms | 80 |
| Participant 6 | 20 | F | AMc | ECoG | 7 (3 RH) | 36 (20 RH) | 36 (20 RH) | 1200 | 64 | 10 | 450ms | 80 |

Additional analyses suggested that although the three observed response types may not be the only response types that exist in the language network, they do capture a substantial amount of the functional heterogeneity in our dataset. First, we repeated the clustering analysis while omitting electrodes below a parametrically varying reliability threshold, and found that the elbow at k=3 became more pronounced (**Figure 2A inset**). Second, when clustering was performed using a larger value of k (e.g., k=10), the profiles of many of the additional clusters resembled the profiles that we discovered when clustering using k=3 (**Figure S2**). And third, responses within a given cluster—especially the more reliable responses—appeared visually similar to the prototypical cluster response profiles, with only a couple of highly reliable responses exhibiting a distinct profile (**Figure S3**).

The average timecourses for the three clusters are shown in **Figure 2E** (see **Figure 2D** for best representative electrodes from each cluster —’medoids’— chosen by the k-medoids algorithm). Cluster 1 (n=92 electrodes; range across participants: 5-34, **Figure S1**) was characterized by a relatively slow increase (build-up) of neural activity across the 8 words in the S condition (a pattern similar to the one reported by Fedorenko et al., 2016; Nelson et al., 2017; Desbordes et al., 2023; Woolnough et al., 2023; but see <u>Discussion</u>), and much lower activity for the W, J, and N conditions, with no difference between the J and N conditions (**Figure 2F**). Cluster 2 (n=67 electrodes; range across participants: 1-21, **Figure S1**) displayed a quicker build-up of neural activity in the S condition that plateaued approximately 3 words into the sentence, a quick build-up of activity in the W condition that began to decay after the third word, and a similar response to the J and N conditions as to the W condition with an overall lower magnitude. Cluster 2 also exhibited ‘locking’ of the neural activity to the onsets of individual words in the S condition. Finally, Cluster 3 (n=18 electrodes; range across participants: 1-7, **Figure S1**) showed no build-up of activity, and was instead characterized by a high degree of locking to the onset of each word or nonword in all conditions. Additionally, the response magnitudes of Cluster 3 were more similar across conditions compared to the other two clusters, although the S>W>J>N pattern was still present (**Figure 2F**).

We then evaluated the stability of these clusters across trials and their robustness to data loss. We found that clusters derived from half of the data (either odd-or even-numbered trials) were significantly more similar to the clusters derived from the full dataset or from the other half of the data than would be expected by chance (ps<0.001, permutation test, <u>Methods</u>, **Figure 3A**). The clusters were also robust to the number of electrodes used: clustering solutions derived from only a subset of the language-responsive electrodes (down to ∼27%, ∼32%, and ∼69% of electrodes for Clusters 1, 2, and 3, respectively) were significantly more similar to the clusters derived from all the electrodes than would be expected by chance (using a threshold of p<0.05, evaluated with a permutation test, <u>Methods</u>, **Figure 3B**).

**Figure 3.**
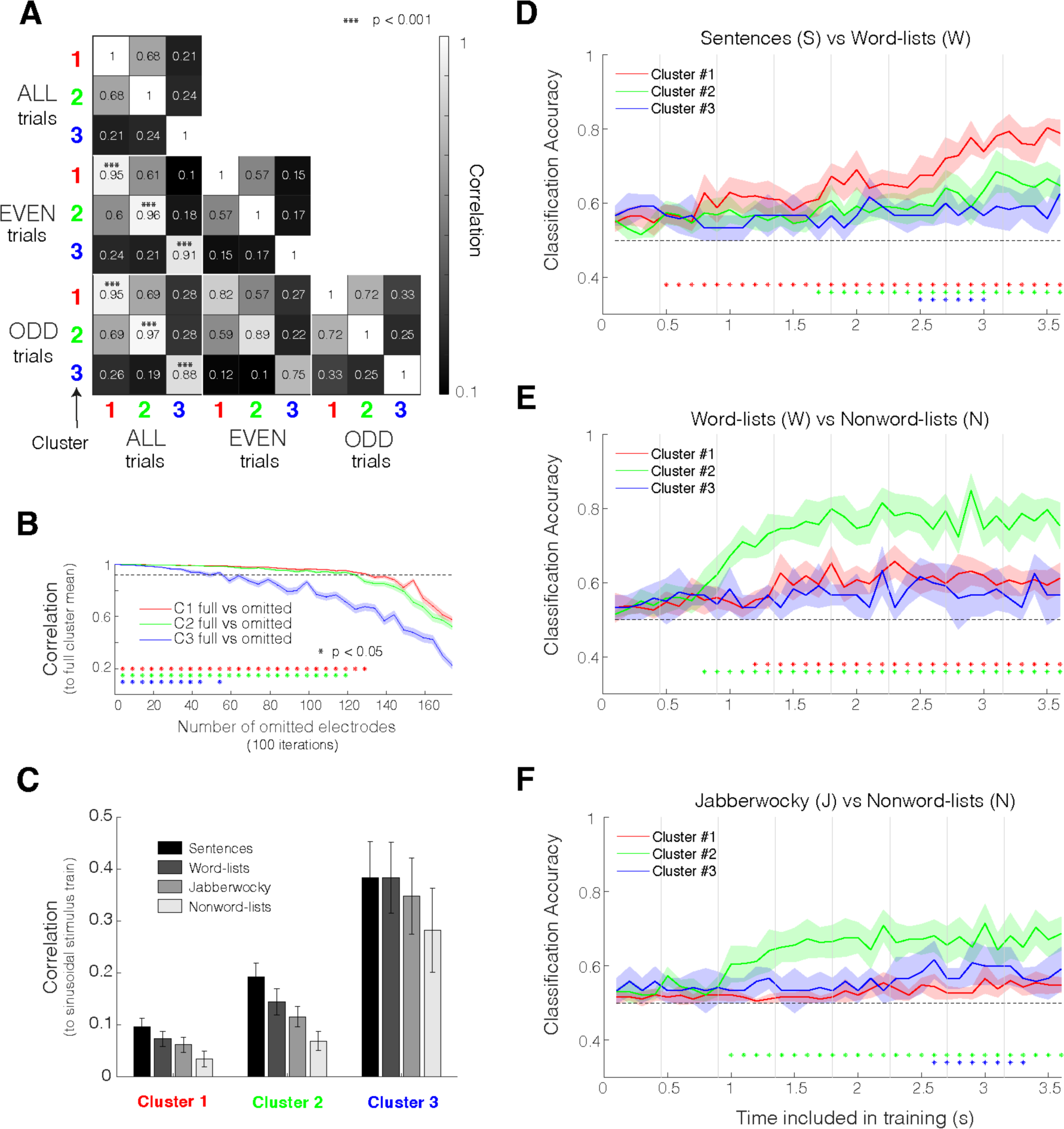
– Evaluation of Dataset 1 clusters. **A**) Comparison of clusters from all trials (top three rows) versus only even (middle three rows) or odd (bottom three rows) trials. Clusters that emerge using only odd or even trials are highly similar to the clusters that emerge when all trials are used (ps<0.001; evaluated with a permutation test; <u>Methods</u>). **B)** Robustness of clusters to electrode omission. Random subsets of electrodes were removed in increments of 5 (<u>Methods</u>). Similarity of cluster centers when all electrodes were used versus when random subsets of electrodes were removed. Stars reflect significant similarity with the full dataset (using a threshold of p<0.05; evaluated with a permutation test; <u>Methods</u>). Shaded regions reflect standard error of the mean over randomly sampled subsets of electrodes. Cluster 3 was driven the most by individual electrodes relative to Clusters 1 and 2. **C)** Correlation of fitted stimulus train with timecourse of electrodes by cluster and by condition (<u>Methods</u>). Error bars reflect standard error of the mean over electrodes. Electrodes in Cluster 3 were the most locked to word/nonword presentation whereas electrodes in Cluster 1 were the least locked to word/nonword presentation. There was a significant main effect for cluster (p<0.05) but not for condition (ANOVA for LME models, <u>Methods</u>, **Table S1A-B**). These qualitative between-condition differences could be due to generally greater engagement of these neural populations with more language-like stimuli. **D-F)** Classifier performance by cluster as a function of the amount of timecourse included in training (<u>Methods</u>). A binary logistic classifier was trained to discriminate the Sentence (S) and Word-list (W) conditions (**D**), Word-list (W) and Nonword-list (N) conditions (**E**), and Jabberwocky (J) and Nonword-list (N) conditions (**F**). Significance stars at the bottom (colored by cluster) reflect discriminability of conditions above chance level (ps<0.05, evaluated as a cluster statistic against a null distribution from permuted labels, <u>Methods</u>). Shaded regions reflect standard error across the 10 folds of the cross-validated classifier.

To further quantify the apparent differences among the three response profiles, we performed two additional analyses. First, we examined how strongly the neural signal exhibited ‘locking’ to individual word/nonword onsets by correlating the observed responses with a fitted sinusoidal function (<u>Methods</u>). This analysis revealed that—consistent with visual examination—electrodes in Cluster 3 showed the strongest degree of stimulus locking, followed by electrodes in Cluster 2, with electrodes in Cluster 1 showing the weakest stimulus-related locking (**Figure 3C**, **Table S1A-B**). And second, we tested how quickly and strongly the S, W, J, and N conditions diverged from one another in each of the profiles. We did this using a binary logistic classifier—trained for each cluster separately—using incrementally more of the timecourse for discrimination (**Figure 3D-F**, <u>Methods</u>). The classification performance (averaged across 10 folds of the cross-validated classifier) revealed that neural populations in Cluster 1 reliably distinguished S from W earlier and more strongly than the neural populations in Clusters 2 and 3. In contrast, neural populations in Cluster 2 reliably distinguished W from N and J from N earlier and more strongly than neural populations in Clusters 1 and 3.

Although the k-medoids clustering algorithm assigns each electrode to one of k discrete clusters, we wanted to additionally evaluate the degree to which single electrode profiles fell between the prototypical cluster response profiles. To do this, we computed the partial correlation of every electrode’s response profile with that of each of the cluster medoids, while controlling for the other two medoids (**Figure S4,** <u>Methods</u>). As shown in **Figure S4B**, many of the electrodes exhibited response profiles that were consistent with only *one* of the prototypical responses. However, a few electrodes, mostly in Clusters 1 and 2, exhibited high partial correlations with another cluster’s medoid (i.e., a “mixed” response profile). Visual inspection of these response profiles (**Figure S4C-D;** osf.io/xfbr8/) revealed that these electrodes displayed a blend of Cluster 1 and Cluster 2 response characteristics. The existence of mixture electrodes primarily between Clusters 1 and 2 is in line with the generally high correlation between their medoids (0.68 between Cluster 1 and 2 medoids versus 0.21 between Cluster 1 and 3, and 0.24 between Cluster 2 and 3; **Figure 3A**).

### 2. Response profiles reflect different sizes of temporal receptive windows

The temporal dynamics of the neural responses across clusters suggested that the observed differences in the response profiles may reflect different ‘temporal receptive windows’ (TRWs). TRWs are a temporal equivalent of spatial receptive fields that corresponds to the amount of the preceding temporal context that affects the processing of the current input (e.g., Hasson et al., 2008, Lerner et al., 2011; Norman-Haignere et al., 2022). In particular, a neural population that only processes information over the span of a single word should exhibit visible evoked responses at the rate of stimulus presentation, reflecting the momentary stimulus-related fluctuations. On the other hand, a neural population that processes information over spans of multiple words should exhibit a response that reflects a more smoothed version of the stimulus train, with no momentary stimulus-related fluctuations. As described in Section 1, the three clusters differed significantly in their degree of locking to the individual word onsets. Cluster 3 showed the strongest locking, followed by Cluster 2, with Cluster 1 showing the weakest amount of locking (**Figure 3C**). Moreover, a neural population that only processes information over the span of ∼a single word (or less) should show little sensitivity to whether nearby words can be composed into phrases. This is the pattern we saw for electrodes in Cluster 3 (**Figure 3D**): these electrodes did not reliably discriminate between the Sentence and Word-list conditions. In contrast, a population that processes information over spans of multiple words should show sensitivity to the composability of nearby words, and thus should strongly discriminate between sentences and word lists. This is the pattern we saw for electrodes in Clusters 1 and 2, with Cluster 1 electrodes showing earlier and stronger discrimination (**Figure 3D**). Note that this greater difference between the Sentence and Word-list conditions for longer-TRW neural populations is presumably due to the fact that linguistic differences between these two conditions become more pronounced for longer word sequences (e.g., see **Figure S5** for evidence from n-gram frequency counts).

To formally test whether the clusters indeed differ in the size of their TRWs, we constructed a toy model wherein we convolved a simplified stimulus train with response functions (gaussian-based ‘kernels’) of varying widths (TRW sizes denoted as σ; **Figure 4A**, see <u>Methods</u> for model assumptions and implementational details). The resulting simulated responses exhibited striking visual similarity to the observed response patterns (**Figure 4A**). We then computed—for every electrode—a correlation between each simulated response and the observed response, and we selected the σ value that yielded the highest correlation (**Figure 4B-C**, <u>Methods</u>). The estimated TRW sizes showed a clear pattern of decrease from Cluster 1 to 2 to 3; the average σ values per cluster were ∼6, ∼4, and ∼1 words for Clusters 1, 2, and 3, respectively (ps<0.0001 comparing TRWs across all pairs of clusters, evaluated with a LME model, <u>Methods</u>, **Figure 4B-C, Table S5**). To evaluate the robustness of this result, we repeated the TRW fitting procedure using other kernel shapes, and confirmed that the relative sizes of the TRWs of the three clusters did not depend on the specific choice of kernel shape (**Figure S6**). Furthermore, the estimated values of σ in number of words (as reported above) appear to be invariant to the stimulus presentation rate, which suggests that the TRW of language-responsive electrodes is information-, not time-, dependent (**Table S6** and **Table S7**). However, this rate-invariance should be investigated further in future work given the small number of participants in each presentation rate group (n=3) and, correspondingly, the low statistical power.

**Figure 4.**
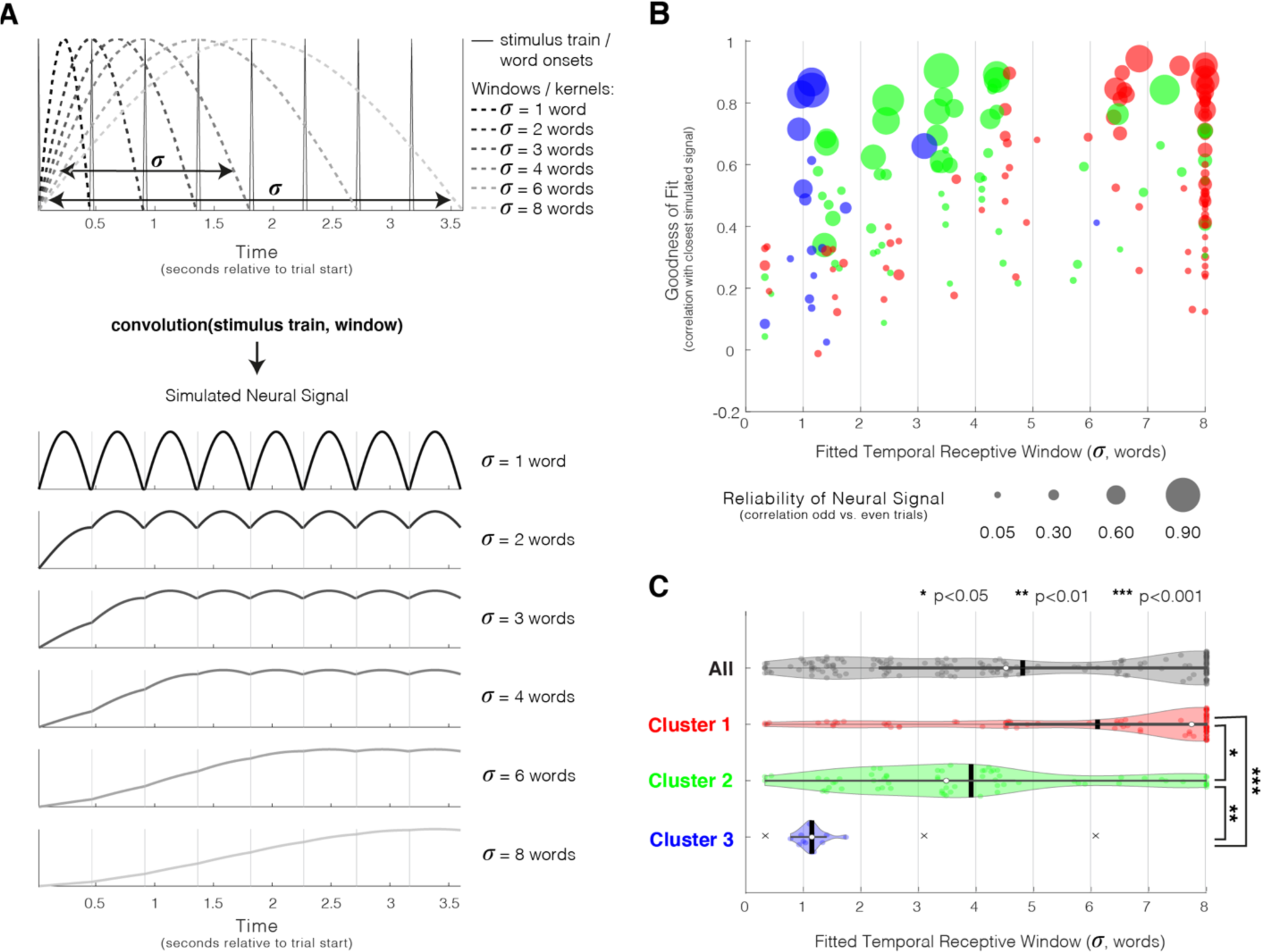
– Estimating the size of the temporal receptive window (TRW) of different electrodes. **A**) A toy model that simulates neural responses to the sentence condition as a convolution of a simplified stimulus train and truncated Gaussian kernels with varying widths. ***Top***: Simplified stimulus train where peaks indicate a word/nonword onset, and sample kernels correspond to varying temporal receptive window sizes (σ). The kernels were constructed from Gaussian curves with a standard deviation of σ/2 truncated at +/− 1 standard deviation (capturing 2/3 of the area under the Gaussian, <u>Methods</u>) and normalized to a minimum of 0 and a maximum of 1. ***Bottom:*** The resulting simulated neural signals for sample kernel widths, normalized to a minimum of 0 and a maximum of 1. **B)** Best TRW fit for all electrodes colored by cluster and sized by the reliability of the neural signal as estimated by correlating responses to odd and even trials (**Figure 1D**). The goodness of fit, or correlation between the simulated and observed neural signal (Sentence condition only), is shown on the y-axis. **C)** Estimated TRW sizes across all electrodes (grey) and per cluster (red, green, and blue). Black vertical lines correspond to the mean window size and the white dots correspond to the median. “x” marks (present in Cluster 3 only) indicate outliers (more than 1.5 interquartile ranges above the upper quartile or less than 1.5 interquartile ranges below the lower quartile). Significance was evaluated with an LME model (<u>Methods</u>, **Table S5**). Together, **B** and **C** show that the clusters varied in the size of their TRWs, from a relatively long TRW (Cluster 1) to a relatively short one (Cluster 3).

### 3. Clusters 1 and 2 are distributed across the language network, whereas cluster 3 exhibits a posterior bias

We tested for differences in the anatomical distribution of the electrodes that belong to the 3 clusters in Dataset 1. We excluded from this analysis right-hemisphere (RH) electrodes because only 4 RH electrodes passed the language selectivity criterion (S>N). We focused on the y (posterior-anterior) and z (inferior-superior) directions in the MNI coordinate space within the left hemisphere. Electrodes in both Clusters 1 and 2 were distributed across the temporal and frontal language regions (**Figure 5**). When examining all electrodes together, or focusing on only the frontal or only the temporal electrodes, the MNI coordinates of electrodes in Clusters 1 and 2 did not significantly differ in either of the two tested directions (ps>0.05, evaluated with a LME model, <u>Methods</u>, **Figure 5C-D**, **Table S2A**). However, when weighting the electrodes by their reliability in the LME model, electrodes in Cluster 1 fell more anteriorly and inferiorly relative to electrodes in Cluster 2 (ps<0.05, evaluated with a LME model, <u>Methods</u>, **Table S2B**). Electrodes in Cluster 3 were located significantly more posteriorly than those in Clusters 1 and 2 (lower y-coordinate values, both Clusters 3 vs. 1 and Clusters 3 vs. 2, ps<0.0001, <u>Methods</u>, **Figure 5C, Table S2A**).

**Figure 5.**
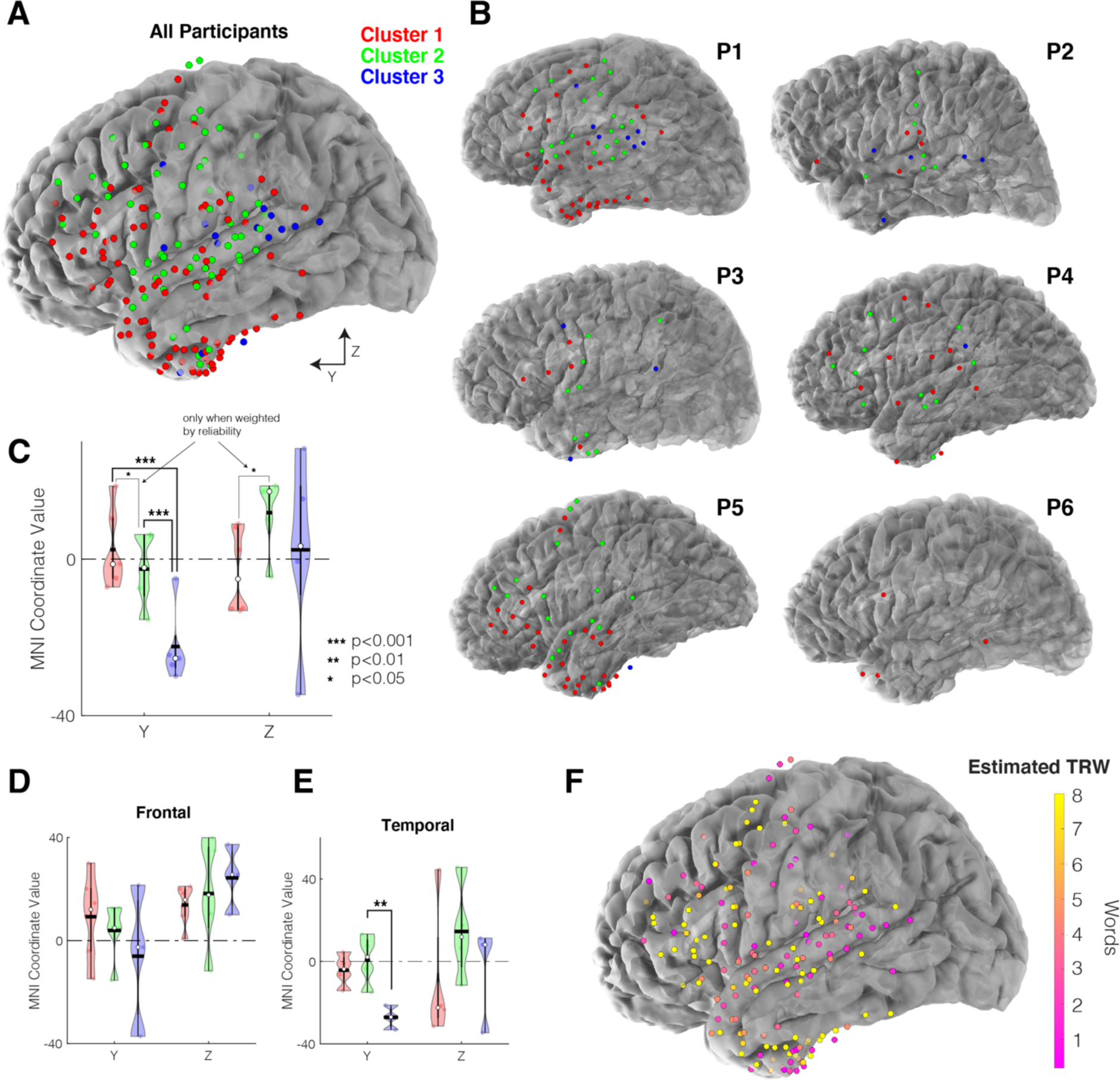
– Anatomical distribution of the clusters in Dataset 1. **A**) Anatomical distribution of language-responsive electrodes in Dataset 1 across all participants in MNI space, colored by cluster. **B)** Anatomical distribution of language-responsive electrodes in participant-specific space. **C-E)** Violin plots of MNI coordinate values for the 3 Clusters, where plotted points represent the mean of all coordinate values for a given participant and cluster. The mean across participants is plotted with a black horizontal line, and the median is shown with a white circle. Significance was evaluated with a LME model (<u>Methods</u>). Cluster 3 exhibited a posterior bias (more negative Y coordinate) relative to Cluster 1 and 2 when modeled using all language electrodes (ps<0.001, **C**). This trend was also evident when examining only the frontal (**D**) or temporal electrodes (**E**) separately, but the difference only reaches significance for the temporal electrodes (p<0.01). **F)** Anatomical distribution of electrodes in Dataset 1 colored by their estimated temporal receptive window (TRW, **Figure 4**). There was a slight trend of increasing TRW size from posterior to anterior regions but with considerable local heterogeneity.

To complement this analysis, we visualized the anatomical distribution of electrodes in two additional ways. First, we visualized all language-responsive electrodes by their partial correlations to each of the cluster medoids (**Figure S4E**). This approach does not enforce a categorical grouping into clusters, potentially allowing for more subtle response gradients. However, this analysis revealed a similar picture: Cluster-1– and Cluster-2-like responses were present throughout frontal and temporal areas, whereas Cluster-3-like responses were localized to the posterior superior temporal gyrus. Second, we examined the distribution of electrodes by their fitted TRW (**Figure 5F**). This visualization exhibited a gross anatomical trend of TRWs increasing from posterior to anterior regions, however, there remained a substantial local mosaic pattern, with long-TRW electrodes present in posterior temporal areas and short-TRW electrodes present in anterior temporal and frontal areas.

### 4. Clusters 1 and 3 replicate in Dataset 2 and Cluster 2 partly replicates

We asked whether the same clusters would emerge in a second, independent dataset with new participants and different linguistic materials (Dataset 2; 16 participants; 362 language-responsive electrodes; mostly depth electrodes, **Figure 6A**, <u>Methods</u>). Participants in Dataset 2 only saw two of the four conditions presented to participants in Dataset 1 (Sentences (S) and Nonword-lists (N), but not Word-lists (W) or Jabberwocky sentences (J)); therefore, we started by re-clustering the electrodes from Dataset 1 using only the responses to the S and N conditions to allow for direct comparisons with Dataset 2.

**Figure 6.**
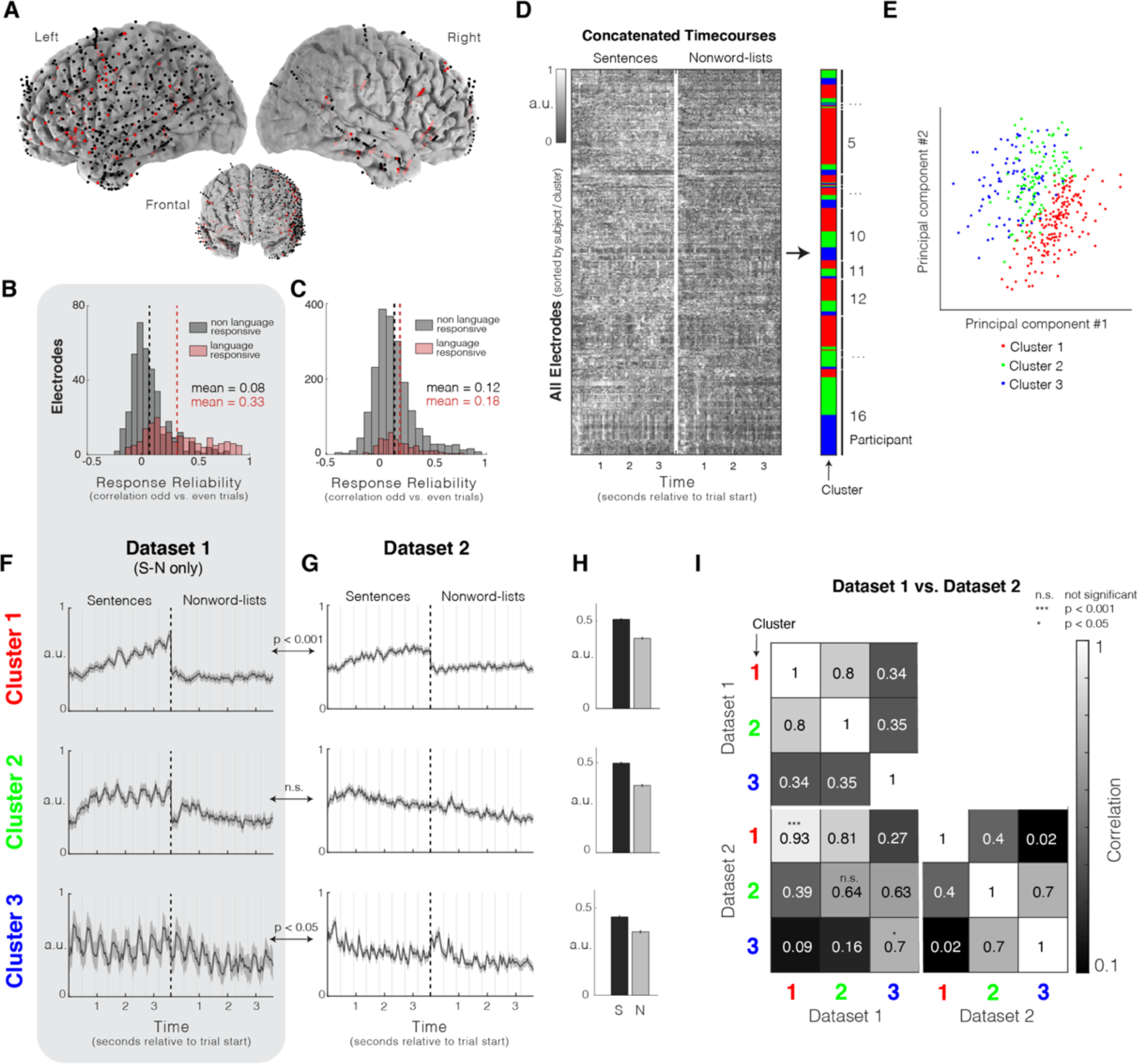
– Dataset 2 k-medoids clustering with k=3. **A**) The locations of language-responsive (n=362, red; <u>Methods</u>) and non-language-responsive (n=2,017, black) electrodes across the sixteen participants in Dataset 2 (both surface and depth electrodes were implanted). Language-responsive electrodes were found across the cortex, in both the left and right hemispheres (**Table 2**). **B and C)** Response reliability as estimated by correlating responses to odd and even trials for language-responsive and non-language-responsive electrodes (as in **Figure 1D**). Language-responsive electrodes exhibit more reliable responses to linguistic stimuli than non-language-responsive electrodes for both Dataset 1 (Sentence and Nonword-list conditions only, **B**) and Dataset 2 (**C**), however, the responses of language electrodes were less reliable in Dataset 2 than Dataset 1. **D)** Clustering mean electrode responses (concatenated responses to sentences and nonword lists) in Dataset 2 using k-medoids (k=3) with a correlation-based distance. Shading of the data matrix reflects normalized high-gamma power (70-150Hz). **E)** Electrodes visualized on their first two principal components, colored by cluster. **F and G)** Average timecourse by cluster from Dataset 1 when using only the Sentence and Nonword-list conditions (**F**; see **Figure S7**) and from Dataset 2 (**G**). Shaded areas around the signal reflect a 99% confidence interval over electrodes. **H)** Mean condition responses by cluster in Dataset 2. Error bars reflect standard error of the mean over electrodes. As with Dataset 1, after averaging across time, response profiles were not as distinct by cluster, underscoring the importance of temporal information in elucidating this grouping of electrodes. **I)** Evaluation of clusters from Dataset 1 (clustering with Sentence and Nonword-list conditions only) against clusters from Dataset 2. Clusters 1 and 3 from Dataset 1 replicated in Dataset 2 (p<0.001 and p=0.023, respectively; permutation test; <u>Methods</u>). Although Cluster 2 demonstrated some qualitative similarity across the two datasets, this similarity was not statistically reliable (p=0.732, permutation test, <u>Methods</u>).

**Table 2.** – Details for Dataset 2. (The data were collected at four sites: Albany Medical Center (Site=AMC), Barnes-Jewish Hospital (Site=BJH), Mayo Clinic Jacksonville (Site=MCJ), and St. Louis Children’s Hospital (Site=SLCH)).

| Participant | Age | Sex | Site | ECoG or sEEG | Language-responsive electrodes (S>N) | Total clean electrodes | Total electrodes | Native sampling freq (Hz) | Elec per amp | Runs | Pres rate (per word) | Trials per cond |
| --- | --- | --- | --- | --- | --- | --- | --- | --- | --- | --- | --- | --- |
| Participant 7 | 51 | M | AMc | ECoG | 14 (7 RH) | 116 (25 RH) | 126 (26 RH) | 1200 | 64 | 3 | 750ms | 48 |
| Participant 8 | 30 | F | AMC | both | 18 (0 RH) | 76 (1 RH) | 92 (3 RH) | 1200 | 64 | 3 | 750ms | 72 |
| Participant 9 | 31 | M | AMC | sEEG | 2 (1 RH) | 90 (44 RH) | 98 (52 RH) | 1200 | 64 | 2 | 600ms | 72 |
| Participant 10 | 59 | F | AMC | sEEG | 2 (0 RH) | 113 (0 RH) | 124 (0 RH) | 1200 | 64 | 2 | 600ms | 72 |
| Participant 11 | 23 | M | AMc | ECoG | 58 (33 RH) | 209 (110 RH) | 216 (110 RH) | 1200 | 64 | 2 | 600ms | 72 |
| Participant 12 | 39 | M | AMC | sEEG | 5 (5 RH) | 112 (112 RH) | 128 (128 RH) | 1200 | 64 | 2 | 600ms | 72 |
| Participant 13 | 29 | M | AMC | ECOG | 9 (0 RH) | 126 (0 RH) | 132 (0 RH) | 1200 | 64 | 2 | 600ms | 72 |
| Participant 14 | 36 | M | AMC | sEEG | 3 (2 RH) | 169 (84 RH) | 184 (90 RH) | 1200 | 64 | 2 | 600ms | 72 |
| Participant 15 | 25 | M | BJH | sEEG | 19 (16 RH) | 183 (93 RH) | 183 (93 RH) | 1000 | 64 | 2 | 600ms | 72 |
| Participant 16 | 38 | M | BJH | sEEG | 49 (15 RH) | 169 (72 RH) | 224 (112 RH) | 1000 | 64 | 2 | 600ms | 72 |
| Participant 17 | 31 | F | BJH | sEEG | 17 (0 RH) | 228 (30 RH) | 228 (30 RH) | 1000 | 64 | 2 | 600ms | 72 |
| Participant 18 | 40 | M | BJH | sEEG | 35 (5 RH) | 137 (11 RH) | 192 (14 RH) | 1000 | 64 | 2 | 600ms | 72 |
| Participant 19 | 66 | M | BJH | sEEG | 32 (1 RH) | 210 (13 RH) | 234 (16 RH) | 2000 | 64 | 2 | 600ms | 72 |
| Participant 20 | 24 | M | BJH | sEEG | 7 (0 RH) | 156 (30 RH) | 218 (30 RH) | 2000 | 64 | 2 | 600ms | 72 |
| Participant 21 | 39 | M | MCJ | sEEG | 11 (1 RH) | 108 (45 RH) | 109 (45 RH) | 1200 | 64 | 1 | 600ms | 36 |
| Participant 22 | 21 | F | SLCH | sEEG | 81 (81 RH) | 176 (176 RH) | 186 (186 RH) | 2000 | 64 | 2 | 600ms | 72 |

The Dataset 1 cluster averages, when only the S and N conditions were used, exhibited a strong qualitative similarity to those of the clusters derived using the data from all four conditions (**Figure S7**). ∼80% of electrodes in Dataset 1 were assigned to the same cluster (‘matched’ to the original clusters by highest correlation). However, Cluster 2 was less robust to electrode loss than Clusters 1 and 3 (compare the green curve in **Figure 3B** to the green curve in **Figure S7G**). This finding suggests that responses to the Word-list and Jabberwocky conditions are especially important for differentiating Cluster 2 from the other response profiles, presumably because these conditions pattern differently for Clusters 1 and 2.

We next clustered the electrodes in Dataset 2 using the same approach as for Dataset 1. The optimal number of clusters in Dataset 2 was k=2 based on the elbow method, and the resulting clusters were visually similar to Clusters 1 and 3 from Dataset 1 (p<0.001 for Cluster 3, p=0.061 for Cluster 1, permutation test, <u>Methods</u>, see OSF osf.io/xfbr8/; note that this permutation test is especially conservative with only two experimental conditions and when k=2). We also performed a version of clustering Dataset 2 enforcing k=3 to test whether a Cluster-2-like response would emerge (**Figure 6**). The same two cluster centers as in the case of k=2 were again apparent and showed reliable similarity to Clusters 1 and 3 in Dataset 1 (p<0.001 and p=0.023, respectively, permutation test, <u>Methods</u>, **Figure 6G, I**). The third cluster qualitatively resembled Cluster 2 from Dataset 1 (**Figure 6G**), but the resemblance was not statistically reliable (p=0.732, permutation test, <u>Methods</u>).

As another, less stringent, test of whether Cluster 2 responses were present in Dataset 2, we assigned each electrode in Dataset 2 to a “group” based on their highest correlation with the average response profiles from Dataset 1, in a “winner-take-all” approach (**Figure S8**). In this approach, a substantial number of electrodes (n=95 of the total of n=362) were assigned to Group 2 (the analog of Cluster 2). This analysis indicates that Cluster-2-like responses are indeed present in Dataset 2, even though they did not reliably emerge through the data-driven clustering approach. The lower robustness of the Cluster-2-like responses in Dataset 2 could be, in part, attributable to the lower split-half reliability of Dataset 2 compared to Dataset 1 (compare **Figure 6B** vs. **6C**), as well as the sparser spatial coverage due to the prevalence of depth electrodes (**Figure 6A**). For completeness, an analysis of the anatomical trends in Dataset 2 is presented in **Figure S9** (**Tables S3** and **S4**).

Finally, we estimated the temporal receptive window (TRW) size (as in Section 2) for each electrode in Dataset 2 (**Figure S10**). Clusters 1 and 3 (the two clusters that consistently replicated from Dataset 1), were best described by TRWs of ∼4.5 and ∼1 words, respectively (**Figure S10A-B**), similar to the TRW sizes observed for those clusters in Dataset 1. The TRW of Cluster 2 did not significantly differ from Cluster 3 when relying on the electrode assignments from the clustering algorithm with k=3 (where Cluster 2 did not replicate, <u>Methods</u>, **Figure 6**, **Figure S10B**, **Table S8**). However, using the winner-take-all approach (where a Cluster-2-like response was “pulled out” into Group 2, **Figure S8**, **Figure S10D**), the TRW of Group 2 was ∼2.1 words, which significantly differed from that of Groups 1 and 3 (ps<0.001 comparing TRWs across all pairs of groups, evaluated using an LME model, <u>Methods</u>, **Figure S10C-D, Table S9**) and was similar to the TRW of Cluster 2 from Dataset 1.

## Discussion

The nature of the neural computations that support our ability to extract meaning from linguistic input remains an important open question in the field of language research. Here, we leveraged the high temporal and spatial resolution of human intracranial recordings to probe the fine temporal dynamics and the spatial distribution of language-responsive neural populations. We uncovered three temporal profiles of response during the processing of sentences and linguistically degraded conditions such as lists of words or nonwords. We suggest that these profiles differ in the size of their temporal receptive window (TRW)—the amount of temporal context that affects the neural processing of the current input. Further, we found that electrodes with distinct response profiles manifest in a scattered spatial distribution across both frontal and temporal cortices. Below, we contextualize these results with respect to prior empirical work and discuss their implications for our understanding of human language processing.

### Three profiles characterize language-responsive neural populations

In the present study, we used a clustering approach in order to group neural populations (as measured by intracranial macroelectrodes; note that when we write that ‘*electrodes’* exhibit a response, we are referring to the *neural populations* that the electrodes are measuring) by their responses to four types of language stimuli: sentences (S), lists of unconnected words (W), Jabberwocky sentences (where content words are replaced with pronounceable nonwords; J), and lists of nonwords (N). We uncovered three dominant response profiles (‘clusters’) that differed in the presence and timing of the increase (build-up) of neural activity over the course of a sentence, the degree of locking to individual word/nonword onsets, and the overall magnitude of response to the linguistically degraded conditions (W, J, and N). Within each cluster, individual electrodes exhibited highly similar responses, with a small number of electrodes displaying a mixed response between Clusters 1 and 2. Finally, we found evidence for each of the three response profiles in an independent dataset that only included two of the four linguistic conditions (Sentences and Nonword-lists), although Clusters 1 and 3 were more robustly replicated. Importantly, because we had restricted our analyses to electrodes that show a functional signature of the language network (a stronger overall response during the processing of structured and meaningful language stimuli—sentences—than during the processing of perceptually similar but meaningless and unstructured stimuli—nonword lists; Fedorenko et al., 2010), these findings provide evidence for ***functional heterogeneity within the language network*** proper, rather than between the language areas and nearby functionally distinct brain regions, like speech areas (e.g., Overath et al., 2015; Keshishian et al., 2023) or higher-level cognitive networks (e.g., Braga et al., 2020; Fedorenko & Blank 2020; Shain, Paunov, Chen et al., 2023; see Fedorenko, Ivanova & Regev, in press, for discussion).

The experimental design adopted in the current study has traditionally been used as a way to tease apart neural responses to word meanings (present in sentences and word lists, but not in Jabberwocky sentences and nonword lists) and syntactic structure (present in sentences and, under some views of syntax, in Jabberwocky sentences, but not in word/nonword lists; Fedorenko et al., 2010, 2012, 2016; for earlier uses of this paradigm, see Mazoyer et al., 1993; Friederici et al., 2000; Humphries et al., 2001; Vandenberghe et al., 2002). As measured with fMRI, all areas of the language network show sensitivity to both word meanings and syntactic structure: the response is strongest to sentences, lower to word lists and Jabberwocky sentences, and lowest to nonword lists (e.g., Fedorenko et al., 2010; Bedny et al., 2011; Shain, Kean et al., in press; Pallier et al., 2011; Desbordes et al., 2023 see Bautista & Wilson, 2016 and Fedorenko et al., 2020 for evidence against the lexical/syntactic dissociation from other paradigms; see Dick et al., 2001 for earlier arguments and evidence). Using a similar design in an intracranial recording study, Fedorenko et al. (2016) replicated this overall pattern of response and also reported a temporal profile—present in a subset of electrodes—whereby high gamma power builds-up across words over the course of a sentence but not in other conditions (replicated by Nelson et al., 2017; Desbordes et al., 2023; Woolnough et al., 2023). They interpreted this build-up effect as indexing the process of constructing a sentence-level meaning.

Here, we investigated the temporal profiles of language-responsive electrodes more comprehensively. By leveraging the fine-grained temporal information in the signal (i.e., considering the full timecourses instead of averaging high gamma power in each word/nonword as in Fedorenko et al., 2016), we found that the build-up effect reported in Fedorenko et al. (2016) represents a mix of functionally distinct populations. The timecourse of response to the Sentence condition in Fedorenko et al. (2016) is most similar to that in Cluster 1 here. However, a reliable sentences > word lists > Jabberwocky sentences > nonword lists profile in Fedorenko et al. (2016) suggests a contribution from Cluster 2 neural populations. As such, our analyses identify two functionally distinct build-up profiles and additionally uncover a third profile, which does not show build-up of activity over time, and we replicated these results in a new, larger dataset with a different set of language materials (Dataset 2). Importantly, here we show that despite strong integration between lexical and syntactic processing, neural populations within the language network *do* differ functionally, although along a different dimension—the temporal scale of information integration.

### The response profiles reflect distinct temporal receptive windows

A temporal receptive window (TRW) denotes the amount of the preceding context that a given neural unit integrates over (e.g., Hasson et al., 2008; Lerner et al., 2011; Norman-Haignere et al., 2022). Previous studies have demonstrated that cortical neural activity is organized into a hierarchy of timescales, wherein information over tens to hundreds of milliseconds is encoded by sensory cortical areas, and information over many seconds is encoded by higher-order areas (Chaudhuri et al., 2015; Runyan et al., 2017; Murray et al., 2014; Chien et al., 2020). Past fMRI studies have shown that the TRW of the language network falls somewhere between a word and a short sentence (e.g., Lerner et al., 2011; Jacoby and Fedorenko, 2020; Blank and Fedorenko, 2020; Jain et al., 2020; Caucheteux et al., 2023; Chang et al., 2022; Shain, Kean et al., in press), although some work has suggested that language regions are, at least to some degree, sensitive to sub-lexical regularities (Bozic et al., 2010; Regev et al., 2024). Using a simple instantiation of an information processing system—with one (interpretable) free parameter: the length of past stimulus context—we estimated the TRW of different language-responsive neural populations. Based on this analysis, we argue that our observed ***response profiles differ in their timescale of information processing***, from sub-lexical units and single words (Cluster 3) to short phrases (Cluster 2) to longer phrases/sentences (Cluster 1).

Do the observed response profiles reflect categorically distinct clusters that integrate information over different timescales, or is the underlying structure of language-selective responses in the brain best described by a continuum of TRWs with no sharp boundaries or groupings of response types? Although we do not rule out the possibility of a TRW continuum, our data are well explained by the grouping of responses into three categories. A few electrodes do exhibit a “mixed” response profile, falling somewhere between the prototypical Cluster 1 and Cluster 2 responses, but this mixing could be due to these electrodes picking up activity of multiple neural populations. Recordings at a higher spatial resolution would be needed to evaluate this possibility (e.g., Paulk et al., 2022; Leonard, Gwilliams et al., 2023). Nevertheless, the current data suggest the existence of neural populations within the language network that are sensitive to information chunks of *distinct and specific size.* This functional organization is presumably driven by the statistics of natural language and is likely critical for efficient extraction of meaning from language (see <u>Future directions</u>).

To estimate the TRW values, we made several simplifying assumptions that can be revisited in future studies. First, we have discussed TRWs in terms of the number of *words*. However, natural languages vary substantially in how they package information into words (Evans & Levinson, 2009) and the processing of a given word is highly dependent on how informative the word is in context (e.g., Shannon, 1949; for behavioral evidence, see Levy, 2008b; Shain et al., 2024). As a result, TRWs may instead be bounded by the number of bits of information. Future work should evaluate multiple accounts of the units in which TRWs are measured. The second simplifying assumption we made was that TRWs are *fixed* in size. Much recent evidence suggests that human comprehension mechanisms can flexibly accommodate corrupt linguistic input, e.g., due to speech errors (e.g., Levy, 2008a; Gibson et al., 2013; Gibson et al., 2017; Keshev & Meltzer-Asscher, 2021; Ryskin et al., 2018, 2021; see Gibson et al., 2019 for a review), which may make it desirable for TRWs to be somewhat adaptable to allow for the possibility of continuously revising one’s interpretation of the input. Future work should seek to understand if and how the TRW of a specific neural population can be affected by linguistic context. And third, the response function (kernel) that we used to generate the simulated signals was intentionally simple and is likely not consistent with the underlying neurophysiology (see <u>Methods</u> for details). A model that is more faithful to neurobiological principles may better capture the observed neural responses and such models should be explored in future work.

Finally, our toy TRW model currently does not take into account the form and content of the stimulus, as it does not use any linguistic information to generate responses. However, responses of neural populations in the language network are highly sensitive to stimulus properties. One key modulator of response strength is how well the stimulus matches natural language statistics, as evidenced by both condition-level effects (e.g., sentences > word lists; Fedorenko et al., 2010) and fine-grained preferences for particular linguistic strings (Tuckute et al., 2024). A more complete model of language processing should therefore include ***both*** “gating” of linguistic input into different lengths of effective input (defined by a neural population’s TRW) ***and*** a scaling of the neural response by the effective input’s probability. This idea—that responses of neural populations in the language network reflect the probability of linguistic inputs at variable context lengths due to their TRW—may explain why the Sentence and Word-list conditions were best discriminated by Cluster 1 populations. In particular, Cluster 1 populations have the longest TRW, and the linguistic difference between sentences and word lists becomes more apparent over longer timescales (as we demonstrated for our stimuli using n-gram probabilities, **Figure S5**). We leave more thorough exploration of stimulus-dependent accounts of the computations carried out by the language network to future work (see <u>Future directions</u>).

### The spatially distributed nature of language processing

There is a long history in language neuroscience of attempts to divide language comprehension into both temporally distinct stages and spatially distinct components. At some level, language comprehension can indeed be broken up across time and space. In particular, clear separation exists between the language-processing system (Fedorenko et al., 2011) and both i) lower-level perceptual areas, and ii) higher-level cognitive areas (see Fedorenko, Ivanova & Regev, in press, for a review). The lower-level perceptual areas, such as the speech perception area (Norman-Haignere et al., 2015; Overath et al., 2015; Keshishian et al. 2023) and the visual word-form area (e.g., Baker et al., 2007; Hamamé et al., 2013; Saygin et al., 2016), process information *earlier* than—and likely provide input to—the language network. And higher-level cognitive areas, such as the areas of the Default network (Buckner & DiNicola, 2019) or the Theory of Mind network (Saxe et al., 2006), process information *later* than—and likely receive input from—the language network. These latter areas plausibly carry out further processing on the meaning representations extracted from language, including connecting those meaning representations across long spans of time (e.g., Lerner et al., 2011; Baldassano et al., 2017; 2018). However, discovering spatial subdivisions *within* the language-selective network proper has proven challenging (e.g., Fedorenko et al., 2010, 2020; Bautista & Wilson, 2016; Blank & Fedorenko, 2020; Shain, Kean et al., in press).

The current work demonstrates that there exist functional differences within the language network, but functionally distinct populations do not seem to exhibit strong spatial clustering and are instead distributed in an interleaved fashion across the language network. The latter explains why most past fMRI work could not reveal this functional heterogeneity (cf. Fedorenko et al., 2012 for implied functional heterogeneity based on multivariate patterns of fMRI response; and see Jain et al., 2020 for evidence of voxel-level heterogeneity with respect to TRWs as discovered in an encoding approach with artificial neural network language models). This architectural design makes it possible for each area of the network to have access to information at different timescales, which likely makes language processing efficient and robust. A clear exception in our data is the concentration of Cluster 3 (shortest-TRW) electrodes in the posterior superior temporal gyrus, which may suggest that this area serves a unique computational role within the language network (see Wilson et al., 2023 and Shain, Kean et al., in press, for other recent evidence of the special role of this area); however, we cannot rule out the possibility that these electrodes are picking up some activity from the nearby speech areas (e.g., Overath et al., 2015). We also acknowledge that a macro-scale organization could become more evident with more participants and a more systematic coverage of the frontal and temporal cortex.

### Future directions

The current findings lay the foundation for several exciting future research avenues. ***First***, the size of a neural unit’s temporal receptive window (TRW) should determine its sensitivity to different linguistic features. As noted above, one limitation of the current investigation is the focus on condition-level differences, rather than trying to explain fine-grained responses to individual linguistic items. The reason for this choice is two-fold. To start, the current linguistic materials were not constructed with the goal of investigating linguistic (e.g., lexical and syntactic) features: in order to make the materials easy to process for diverse populations, the sentences were constructed to be short and to use common structures and words, which limits the range of variability to be explored. And additionally, we did not observe reliable stimulus-related activity (beyond the level of conditions; see OSF osf.io/xfbr8/). However, the TRW-based framework makes clear predictions that can be evaluated in future work. For example, short-TRW populations should show greater sensitivity to lexical features, such as word frequencies, whereas longer-TRW populations should be more sensitive to linguistic features at longer timescales, such as higher-order n-gram frequencies and syntactic-structure-related features. Because many linguistic features are strongly inter-correlated in naturalistic language materials (e.g., Piantadosi et al., 2011; Shain, Blank et al., 2020; Shain et al., 2022; see OSF osf.io/xfbr8/ for evidence of inter-correlation of linguistic features in the current stimuli), evaluating these predictions will require constructing materials that are specifically designed to best dissociate different linguistic dimensions.

***Second***, artificial neural network (ANN) language models—which have proven to be powerful tools for understanding the human language system (Toneva & Wehbe, 2019; Jain et al., 2020; Schrimpf et al., 2021; Goldstein et al., 2022; Caucheteux & King, 2022; see Tuckute et al., in press, for a review)—could be leveraged to gain insights into the constraints on the language processing architecture. For example, do successful language architectures require particular proportions of units with different TRWs or particular distributions of such units within and/or across model layers? In Dataset 1, we found the fewest electrodes belonging to Cluster 3 (shortest TRW), more electrodes belonging to Cluster 2 (intermediate TRW), and the majority of electrodes belonging to Cluster 1 (longest TRW). These proportions align with the idea that compositional semantic space is highly multi-dimensional, but word-form information can be represented in a relatively low-dimensional space (e.g., Mollica and Piantadosi, 2019). However, the proportions can also be affected by biases in where intracranial electrodes tend to be implanted, so investigating these questions in ANNs, where we can probe all units in the network and have the freedom to alter the architecture in various ways, may yield insights that cannot be gained from human brains at least with the current experimental tools available.

And ***third***, we have here focused on language comprehension. However, the same language network also supports language production (Awad et al., 2007; Menenti et al. 2011; Segaert et al. 2012; Silbert et al., 2014; Giglio et al., 2022; Hu, Small et al., 2022). Whether the TRW-based organization discovered here in a language comprehension task applies to language production— given that utterance planning is known to unfold at multiple scales (e.g., Lee et al., 2013)— remains to be determined.

In conclusion, across two intracranial-recording datasets, we here demonstrate the existence of functionally distinct neural populations within the fronto-temporal language-selective network proper, opening the door to investigations of how these populations work together to accomplish the incredible feats of language comprehension and production.

## Methods

### Participants

*Dataset 1* (also used in Fedorenko et al., 2016): Electrophysiological data were recorded from intracranial electrodes in 6 participants (5 female, aged 18–29 years) with intractable epilepsy. These participants underwent temporary implantation of subdural electrode arrays at Albany Medical Center to localize the epileptogenic zones and to delineate it from eloquent cortical areas before brain resection. All participants gave informed written consent to participate in the study, which was approved by the Institutional Review Board of Albany Medical Center. One further participant was tested but excluded from analyses because of difficulties in performing the task (i.e., pressing multiple keys, looking away from the screen) during the first five runs. After the first five runs, the participant required a long break during which a seizure occurred.

*Dataset 2:* Electrophysiological data were recorded from intracranial electrodes in 16 participants (4 female, aged 21-66 years) with intractable epilepsy. These participants underwent temporary implantation of subdural electrode arrays and depth electrodes to localize the epileptogenic zones before brain resection at one of four sites: Albany Medical Center (AMC), Barnes-Jewish Hospital (BJH), Mayo Clinic Jacksonville (MCJ), and St. Louis Children’s Hospital (SLCH). All participants gave informed written consent to participate in the study, which was approved by the Institutional Review Board at each relevant site. Two further participants were tested but excluded from analyses due to the lack of any language-responsive electrodes (see <u>Language-</u> <u>Responsive Electrode Selection</u>).

### Data Collection

*Dataset 1:* The implanted electrode grids consisted of platinum-iridium electrodes that were 4 mm in diameter (2.3–3 mm exposed) and spaced with an inter-electrode distance of 0.6 or 1 cm. The total numbers of implanted grid/strip electrodes were 120, 128, 98, 134, 98, and 36 for the six participants, respectively (**Table 1**). Electrodes were implanted in the left hemisphere for all participants except P6, who had bilateral coverage (16 left hemisphere electrodes). Signals were digitized at 1,200 Hz.

*Dataset 2:* The implanted electrode grids and depth electrodes consisted of platinum-iridium electrodes. Implanted grid contacts were spaced at 0.6 or 1cm (2.3–3 mm exposed), while SEEG leads were spaced 3.5 – 5 mm depending on the trajectory length, with 2 mm exposed. The total numbers of implanted electrodes by participant can be found in **Table 2** (average=167 electrodes; st. dev.=51; range 92-234), along with the frequencies at which the signals were digitized. Electrodes were implanted in only the left hemisphere for 2 participants, in only the right hemisphere for 2 participants, and bilaterally for 12 participants (**Table 2**). All participants, regardless of the lateralization of their coverage, were included in all analyses.

For both datasets, recordings were synchronized with stimulus presentation and stored using the BCI2000 software platform (Schalk et al., 2004).

### Cortical Mapping

Electrode locations were obtained from post-implantation computerized tomography (CT) imaging and co-registered with the 3D surface model of each participant’s cortex—created from the preoperative anatomical MRI image—using the VERA software suite (Adamek et al., 2022). Electrode locations were then transformed to MNI space within VERA via nonlinear co-registration of the subjects’ skull-stripped anatomical scan and the skull-stripped MNI152 Freesurfer template using ANTs (Avants et al., 2008).

### Preprocessing and Extraction of Signal Envelope

Neural recordings were collected and saved in separate data files by run (see Experiment and **Tables 1-2**), and all preprocessing procedures were applied *within* data files to avoid inducing artifacts around recording breaks.

First, the ECoG/sEEG recordings were high-pass filtered at the frequency of 0.5 Hz, and line noise was removed using IIR notch filters at 60, 120, 180, and 240 Hz. The following electrodes were excluded from analysis: a) ground, b) reference, and c) those that were not ECoG or sEEG contacts (e.g., microphone electrodes, trigger electrodes, scalp electroencephalography (EEG) electrodes, EKG electrodes), as well as d) those with significant line noise, defined as electrodes with line noise greater than 5 standard deviations above other electrodes, e) those with large artifacts identified through visual inspection, and, for all but four participants, f) those that had a significant number of interictal discharges identified using an automated procedure (Janca et al., 2015). (For 4 participants—P3 in Dataset 1 and P15, P17, and P21 in Dataset 2—electrodes that were identified as having a significant number of interictal discharges were not excluded from analyses because more than 1/3 of each of these participants’ electrodes fit this criterion.) These exclusion criteria left 108, 115, 92, 106, 93, and 36 electrodes for analysis for the 6 participants in Dataset 1 (**Table 1**) and between 76 and 228 electrodes for the 16 participants in Dataset 2 (**Table 2**).

Next, the common average reference (from all electrodes connected to the same amplifier) was removed for each timepoint separately. The signal in the high gamma frequency band (70 Hz– 150 Hz) was then extracted by taking the absolute value of the Hilbert transform of the signal extracted from 8 gaussian filters (center frequencies: 73, 79.5, 87.8, 96.9, 107, 118.1, 130.4, and 144; standard deviations (std): 4.68, 4.92, 5.17, 5.43, 5.7, 5.99, 6.3, and 6.62, respectively, as in e.g., Dichter et al., 2018). The resulting envelopes from each of the Gaussian filters were averaged into one high gamma envelope. We focus on the high gamma frequency range because this component of the signal has been shown to track neural activity most closely (e.g., Janca et al., 2015). Linear interpolation was used to remove data points whose magnitude was more than 5 times the 90^th^ percentile of all magnitudes (Norman-Haignere et al., 2022), and we downsampled the signal by a factor of 4. For all data analysis basic Matlab (version 2021a) functions were used.

Finally, the data were z-scored and normalized to a min/max value of 0/1 to allow for comparisons across electrodes, and the signal was downsampled further to 60 Hz (regardless of the participant’s native sampling frequency) to reduce noise and standardize the sampling frequency across participants. For the participants who performed a slower version of the paradigm (e.g., words presented for 700 ms each; see <u>Experiment</u>), the signal was time-warped to a faster rate (words presented for 450 ms each) so that timecourses could be compared across subjects. This time-warping was done by resampling (Matlab procedure *resample*).

### Experiment

*Dataset 1:* In an event-related design, participants read sentences, lists of words, Jabberwocky sentences, and lists of nonwords. All stimuli were eight words/nonwords long. The materials were adapted from Fedorenko et al. (2010; Experiment 2) and the full details of stimulus construction are described there. In short, sentences were manually constructed to cover a wide range of topics using various syntactic structures. Sentences were intended to be easily read, to fit participants with diverse clinical conditions and only included mono– and bi-syllabic words. The full list of materials is available at OSF (https://osf.io/xfbr8/). The word lists were created by scrambling the words from the sentences. Jabberwocky sentences were created from the sentences by removing content words (e.g., nouns, verbs, etc.), but leaving the syntactic frame, consisting of function words (e.g., articles, conjunctions, prepositions, pronouns, etc.), intact. Content words were replaced with other pronounceable nonwords, matched for length (in syllables). Lastly, the nonword lists were generated from scrambling the words/nonwords from the Jabberwocky condition. Originally, a set of 160 items per each condition were created and here, 80 or 60 items of those were used (depending on stimulus presentation rate, as detailed below).

Each event (trial) consisted of eight words/nonwords, presented one at a time at the center of the screen. At the end of each sequence, a memory probe was presented (a word in the Sentence and Word-list conditions, and a nonword in the Jabberwocky and Nonword-list conditions) and participants had to decide whether the probe appeared in the preceding sequence by pressing one of two buttons. Two different presentation rates were used: P1, P5, and P6 viewed each word/nonword for 450 ms (fast-timing), and P2, P3, and P4 viewed each word/nonword for 700 ms (slow-timing). The presentation speed was determined before the experiment based on the participant’s preference. After the last word/nonword in the sequence, a fixation cross was presented for 250 ms, followed by the probe item (1,400-ms fast-timing, 1,900 ms slow-timing), and a post-probe fixation (250 ms). Behavioral responses were continually recorded, but only responses 1 second before and 2 seconds after the probe were considered for calculating behavioral performance (**Table 3**). Participants performed best on the sentence trials and worst on the nonword list trials, with an average accuracy across all conditions of 81.01% (**Table 3**). After each trial, a fixation cross was presented for a variable amount of time, semi-randomly selected from a range of durations from 0 to 11,000 ms, to obtain a low-level baseline for neural activity.

**Table 3.**
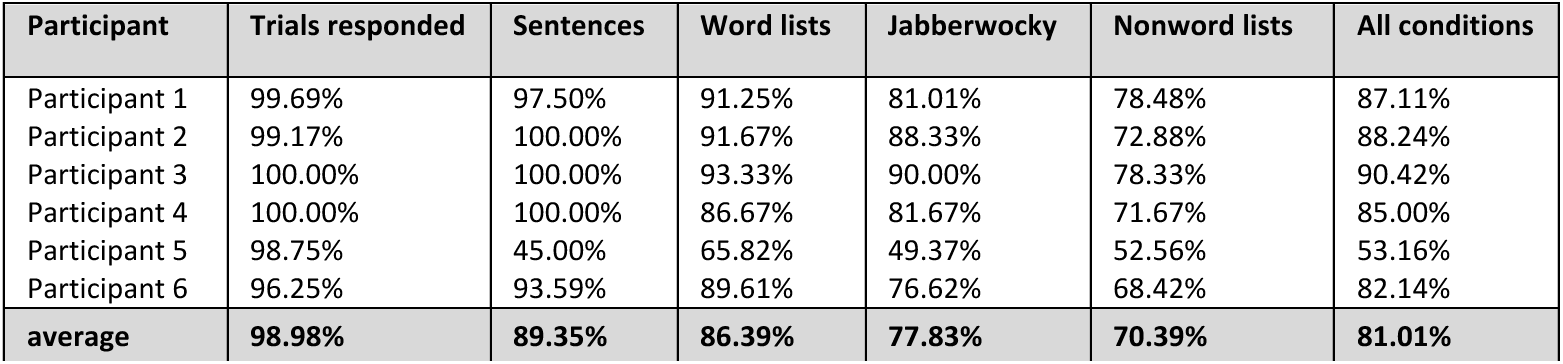
– Behavioral results for Dataset 1. Percentage of trials where participants in Dataset 1 responded and their accuracy on completed trials.

| Participant | Trials responded | Sentences | Word lists | Jabberwocky | Nonword lists | All conditions |
| --- | --- | --- | --- | --- | --- | --- |
| Participant 1 | 99.69% | 97.50% | 91.25% | 81.01% | 78.48% | 87.11% |
| Participant 2 | 99.17% | 100.00% | 91.67% | 88.33% | 72.88% | 88.24% |
| Participant 3 | 100.00% | 100.00% | 93.33% | 90.00% | 78.33% | 90.42% |
| Participant 4 | 100.00% | 100.00% | 86.67% | 81.67% | 71.67% | 85.00% |
| Participant 5 | 98.75% | 45.00% | 65.82% | 49.37% | 52.56% | 53.16% |
| Participant 6 | 96.25% | 93.59% | 89.61% | 76.62% | 68.42% | 82.14% |
| average | 98.98% | 89.35% | 86.39% | 77.83% | 70.39% | 81.01% |

Trials were grouped into runs to give participants short breaks throughout the experiment. In the fast-timing version of the experiment, each run included eight trials per condition and lasted 220 s, and in the slow-timing version, each run included six trials per condition and lasted 264 s. The total amount of intertrial fixation in each run was 44 s for the fast-timing version and 72 s for the slow-timing version. All participants completed 10 runs of the experiment, for a total of 80 trials per condition in the fast-timing version and 60 trials per condition in the slow-timing version. P1 was accidentally shown one run twice, and consequently saw only 9 unique runs for a total of 72 trials per condition (as they opted for the fast presentation rate).

*Dataset 2:* In an event-related design that was similar to the one used in Dataset 1, participants read sentences and lists of nonwords (the other two conditions—lists of words and Jabberwocky sentences—were not included). The materials were adapted from a version of the language localizer in use in the Fedorenko lab (e.g., Lipkin et al., 2022). The sentences came from a language corpus (Brown corpus; Kučera et al., 1967) where we searched for 12-word long sentences and chose a diverse set among those. The nonwords were created using the Wuggy software to match to the words from the sentences on low-level phonology.

Each event (trial) consisted of twelve words/nonwords, presented one at a time at the center of the screen. At the end of each sequence, a memory probe was presented (a word in the Sentence condition and a nonword in the Nonword-list condition) and participants had to decide whether the probe appeared in the preceding sequence by pressing one of two buttons. Two presentation rates were used: 600 ms per word/nonword (medium-timing) and 750 ms per word/nonword (slow-timing; see **Table 2** for a description of the presentation rates by participant). The presentation speed was determined before the experiment based on the participant’s preference. After the last word/nonword in the sequence, a fixation cross was presented for 400 ms, followed by the probe item (1,000 ms for both fast– and slow-timing), and a post-probe fixation (600 ms). Behavioral responses were continually recorded, but only responses 1 second before and 2 seconds after the probe were considered for calculating behavioral performance (**Table 4**). As in Dataset 1, participants performed best on the sentence trials and worse on the nonword-list trials. However, in this sample of participants there was substantial individual variability in the consistency and accuracy of responses (**Table 4**). On average participants provided a correct response 68.57% of the time (**Table 4**). After each trial, a fixation cross was presented for a variable amount of time, semi-randomly selected from a range of durations from 0 to 6,000 ms.

**Table 4.** – Behavioral results for Dataset 2. Percentage of trials where participants in Dataset 2 responded and their accuracy on completed trials.

| Participant | Trials responded | Sentences | Nonword lists | All conditions |
| --- | --- | --- | --- | --- |
| Participant 7 | 79.17% | 70.00% | 75.00% | 72.37% |
| Participant 8 | 95.83% | 88.41% | 81.16% | 84.78% |
| Participant 9 | 45.83% | 50.00% | 57.69% | 53.03% |
| Participant 10 | 98.61% | 94.44% | 65.71% | 80.28% |
| Participant 11 | 16.67% | 40.00% | 44.44% | 41.67% |
| Participant 12 | 99.31% | 93.06% | 64.79% | 79.02% |
| Participant 13 | 86.81% | 83.87% | 76.19% | 80.00% |
| Participant 14 | 99.31% | 97.18% | 79.17% | 88.11% |
| Participant 15 | 95.14% | 71.01% | 55.88% | 63.50% |
| Participant 16 | 0.69% | 0.00% | 0.00% | 0.00% |
| Participant 17 | 83.33% | 95.16% | 79.31% | 87.50% |
| Participant 18 | 90.97% | 92.65% | 76.19% | 84.73% |
| Participant 19 | 100.00% | 94.44% | 83.33% | 88.89% |
| Participant 20 | 34.72% | 57.14% | 36.36% | 48.00% |
| Participant 21 | 52.78% | 46.15% | 64.00% | 57.89% |
| Participant 22 | 98.61% | 91.55% | 83.10% | 87.32% |
| average | 73.61% | 72.82% | 63.90% | 68.57% |

Trials were grouped into runs to give participants short breaks throughout the experiment. In the medium-timing version of the experiment, each run included 36 trials per condition and lasted ∼898 s, and in the slow-timing version, each run included 24 trials per condition and lasted 692 s. The total amount of intertrial fixation in each run was 216 s for the medium-timing version and 144 s for the slowest-timing version. One participant (P7) saw a modified slow-timing version of the paradigm where only 48 of the full 72 items per condition were shown. 13 participants completed 2 runs of the experiment (all saw the medium-timing version, 72 trials per condition), 2 participants completed 3 runs of the experiment (one saw the slow-timing version, 72 trials per condition; and the other saw the modified slow-timing version, 48 trials per condition), and 1 participant completed 1 run of the experiment (medium-timing version, 36 trials per condition, **Table 2**).

For all clustering analyses, only the first eight words/nonwords of the stimulus were used to ensure that the length of the timecourses being analyzed was the same across Dataset 1 and 2.

### Language-Responsive Electrode Selection

In both datasets, we identified language-responsive electrodes as electrodes that respond significantly more (on average, across trials) to sentences (the S condition) than to perceptually similar but linguistically uninformative (i.e., meaningless and unstructured) nonword sequences (the N condition). First, the envelope of the high-gamma signal was averaged across word/nonword positions (8 positions in the experiment used in Dataset 1, and 12 positions in the experiment used in Dataset 2) to construct an ‘observed’ response vector for each electrode (1 x nTrialsS + nTrialsN; the number of trials, across the S and N conditions, varied by participant between 72 and 160). The observed response vector was then correlated (using Spearman’s correlation) with an ‘idealized’ language response vector, where sentence trials were assigned a value of 1 and nonword trials—a value of –1. The values in the ideal response vector were then randomly permuted without replacement and a new correlation was computed. This process was repeated 10,000 times, for each electrode separately, to construct a null distribution (with shuffled labels) relative to which the true correlation between the observed values and the ‘idealized’ values could be evaluated. Electrodes were determined to be language-responsive if the observed vs. idealized correlation was greater than 95% of the correlations computed using the permuted idealized response vectors (equivalent to p < 0.05). (We chose a liberal significance threshold in order to maximize the number of electrodes to be included in the critical analyses, and to increase the chances of discovering distinct response profiles.) The majority of the language-responsive electrodes (98.3% in Dataset 1, 53.9% in Dataset 2) fell in the left hemisphere, but we use electrodes across both hemispheres in all analyses (see e.g., Lipkin et al., 2022 for evidence of a robust right-hemisphere component of the language network in a dataset of >800 participants).

### Clustering analysis

Using Dataset 1 (n=6 participants, m=177 language-responsive electrodes), we created a single timecourse per electrode by concatenating the average timecourses across the four conditions (sentences (S), word lists (W), Jabberwocky sentences (J), nonword lists (N)). All the timepoints of the concatenated timecourses (864 data points: 60 Hz * 4 conditions * 3.60 seconds per trial after resampling) served as input to a k-medoids clustering algorithm (Kaufman & Rousseuw, 1990). K-medoids is a clustering technique that divides data points—electrodes in our case—into k groups, where k is predetermined. The algorithm attempts to minimize the distances between each electrode and the cluster center, where cluster centers are represented by ‘medoids’ (exemplar electrodes selected by the algorithm) and the distance metric is correlation-based. K-medoids clustering was chosen over the more commonly used k-means clustering to allow for the use of a correlation-based distance metric as we were most interested in the shape of the timecourses over their scale which can vary due to cognitively irrelevant physiological differences (but see **Figure S1** for evidence that similar clusters emerge with a k-means clustering algorithm using a Euclidean distance).

### Optimal number of clusters

To determine the optimal number of clusters, we used the “elbow” method (e.g., Rokach and Maimon, 2005) which searches for the value of k above which the increase in explained variance becomes more moderate. For each k (between 2 and 10), k-medoids clustering was performed, and explained variance was computed as the sum of the correlation-based distances of all the electrodes to their assigned cluster center and normalized to the sum of the distances for k=1 (equivalent to the variance of the full dataset). This explained variance was plotted against k and the “elbow” was determined as the point after which the derivative became more moderate. We plot the derivative of this curve as well for easier inspection of the transition point. We also repeat the elbow method while enforcing a parametrically sampled reliability threshold (from 0.1 to 0.4 in increments of 0.1) to further examine our choice of k. If the chosen k does, in fact, appropriately describe the data, we would expect the strength of the elbow (that is, the drop in explained variance for k+1) to increase.

### Partial correlation of individual electrodes with each of the cluster medoids

To evaluate the extent to which the observed responses can be attributed to a single profile, we computed partial correlations (Fisher, 1924) of every electrode’s mean timecourse with that of each of the cluster medoids, while controlling for the other two cluster medoids. For instance, take r*_s_*_1*c*1,*c*2*c*3_ as the partial correlation between a signal *s1* and Cluster 1 medoid *C1*, while controlling for the Cluster 2 medoid *C2* and Cluster 3 medoid *C3*. r*_s_*_1*c*1,*c*2*c*3_ can be computed by, i) performing a multiple regression analysis with *s1* as the dependent variable and *C2* and *C3* as the independent variables, obtaining the residual *e1*; ii) performing a multiple regression analysis with *C1* as the dependent variable and *C2* and *C3* as the independent variable, obtaining the residual *e2*; and iii.) calculating the correlation coefficient between the residuals *e1* and *e2*. This is the partial correlation r*_s_*_1*c*1,*c*2*c*3_. The analysis was performed using the Matlab *partialcorr* function.

### Cluster stability across trials

We evaluated the stability of the clustering solution by performing the same clustering procedure as described above (<u>Clustering analysis</u>) on half the trials. To evaluate the similarity of the clusters derived based on only half of the trials to the clusters derived based on all trials, we first had to determine how clusters correspond between any two solutions. In particular, given that the specific order of the clusters that the k-medoids algorithm produces depends on the (stochastic) choice of initial cluster medoids, the electrodes that make up Cluster 1 in one solution may be labeled as Cluster 2 in another solution. To determine cluster correspondence across solutions, we matched the cluster centers (computed here as the average timecourse of all electrodes in a given cluster) from a solution based on half of the trials to the most highly correlated cluster centers from the solution based on all trials.

We then conducted a permutation analysis to statistically compare the clustering solutions. This was done separately for each of the two halves of the data (odd– and even-numbered subsets of trials). Under the null hypothesis, no distinct response profiles should be detectable in the data, and consequently, responses in one electrode should be interchangeable with responses in another electrode. Using half of the data, we shuffled average responses across electrodes (within each condition separately, thus disrupting the relationship between the conditions for a given electrode while leaving the distribution of within-condition average responses intact), re-clustered the electrodes into 3 clusters, and then correlated the resulting cluster centers to the ‘corresponding’ cluster centers from the full dataset. This permutation test was determined to be more conservative than shuffling individual trials across electrodes (within each condition separately). However, comparisons remained significant when shuffling individual trials. We repeated this process 1,000 times to construct a null distribution of the correlations for each of the 3 clusters. These distributions were used to calculate the probability that the correlation between the clusters across the two solutions using the actual, non-permuted data was higher than would be expected by chance.

### Cluster robustness to data loss

We evaluated the robustness of the clustering solution to loss of electrodes to ensure that the solution was not driven by particular electrodes or participants.

To evaluate the similarity of the clusters derived based on only a subset of language-responsive electrodes to the clusters derived based on all electrodes, we progressively removed electrodes from the full set (n=177) until only 3 electrodes remained (the minimal number of electrodes required to split the data into 3 clusters) in increments of 5. Each subset of electrodes was clustered into 3 clusters, and the cluster centers were correlated with the corresponding cluster centers (see section <u>Cluster stability across trials</u> above) from the full set of electrodes. For each subset of electrodes, we repeated this process 100 times, omitting a different random set of n electrodes with replacement, and computed the average correlation across repetitions.

To statistically evaluate whether the clustering solutions with only a subset of electrodes were more similar to the solution on the full set of electrodes on average (across the 100 repetitions at each subset size) than would be expected by chance, we conducted a permutation analysis like the one described in <u>Cluster stability across trials</u>. In particular, using the full dataset, we shuffled average responses across electrodes (within each condition separately), re-clustered the electrodes into 3 clusters, and then correlated the resulting cluster averages to cluster averages from the actual, non-shuffled data. We repeated this process 1,000 times to construct a null distribution of the correlations for each of the 3 clusters. These distributions were used to calculate the probability that the correlation between the clusters across the two solutions using the actual, non-permuted data (a solution on all the electrodes and a solution on a subset of the electrodes) was higher than would be expected by chance. To err on the conservative side, we chose the null distribution for the cluster with the highest average correlation in the permuted version of the data. For each subset of electrodes, if the average correlation (across the 100 repetitions) fell below the 95^th^ percentile of the null distribution, this was taken to suggest that the clustering solution based on a subset of the electrodes was no longer more correlated to the solution on the full set of electrodes than would be expected by chance.

### Electrode locking to onsets of individual word/nonwords

To estimate the degree of stimulus locking for each electrode and condition separately, we fitted a sinusoidal function that represented the stimulus train to the mean of the odd trials and then computed the Pearson correlation between the fitted sinusoidal function and the mean of the even trials. For the sinusoidal function fitting, we assumed that the frequency of the sinusoidal function was the frequency of stimulus presentation and we fitted the phase, amplitude and offset of the sinusoid by searching parameter combinations that minimized the sum of squared differences between the estimated sinusoidal function and the data. Cross-validation (fitting on odd trials and computing the correlation on even trials) ensured non-circularity. To statistically quantify differences in the degree of stimulus locking between the clusters and among the conditions, we ran a linear mixed-effects (LME, using the Matlab procedure *fitlme*) model regressing the locking values of all electrodes and conditions on the fixed effects categorical variable of *cluster* (with 3 levels for Cluster 1, 2 or 3 according to which cluster each electrode was assigned to) and *condition* (with 4 levels for conditions S, W, J, N), both grouped by the random effects variable of *participant*, as well as a fixed interaction term between *cluster* and *condition*:

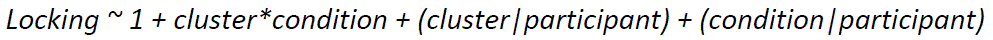

An ANOVA test for LME was used to determine the main effects of *cluster* and *condition* and their interaction. Pairwise comparisons of all 3 clusters and 4 conditions as well as interactions between all (cluster, condition) pairs were extracted from the model estimates.

### Electrode discrimination between conditions

To examine the *timecourse of condition divergence*, as quantified by the electrodes’ ability to linearly discriminate between the magnitudes of pairs of conditions. We focused on condition pairs that critically differ in their engagement of particular linguistic processes: conditions S and W, which differ in whether they engage combinatorial (syntactic and semantic) processing (S=yes, W=no), conditions W and N, which differ in whether they engage word meaning processing (W=yes, N=no), and conditions J and N, which differ in whether they engage syntactic processing (J=yes, N=no). This analysis tests how early the relevant pair of conditions reliably diverge and the strength of that divergence. For every electrode, the mean response to the three conditions of interest (S, W, and N) was averaged across 100 ms bins with a 100 ms sliding window. For each cluster separately, a binary logistic classifier (selected from the best of 20 random instantiations; performed using the Matlab *fitclinear* function) was trained (to discriminate S from W, W from N, or J from N) at each time bin using the binned neural signal up to, and including, that time bin. Each classifier was trained using 10-fold cross validation (train on 90% of the data and test using the remaining 10%, repeated for 10 splits of the data such that every observation was included in the test set exactly once). The predicted and actual conditions across all folds were used to calculate accuracy (the percent of mean responses from all electrodes in a particular cluster correctly classified as S/W, W/N, or J/N). The performance of the model at a given time bin was statistically evaluated using a cluster permutation test to control for multiple comparisons and account for the autocorrelation structure of the signals (Stelzer et al., 2013; Maris & Oostenveld, 2007). This was done by shuffling the condition labels 1000 times for each time bin to simulate surrogate data. For each surrogate data repetition, we computed the sum of consecutive t-values that passed some arbitrary t-value threshold, referred to as the t-sum statistics. We chose a t-value threshold corresponding to an alpha level of 0.05. Using the t-sum values from the 1000 permutations, we constructed a null distribution for this t-sum statistic, and then compared it to the same t-sum statistic computed from the real data to assess significance.

### Computing Ngram frequencies of sentence and nonword stimuli

N-gram frequencies were extracted from the Google n-gram online platform (https://books.google.com/ngrams/), averaging across Google books corpora between the years 2010 and 2020. For each individual word, n-gram frequency for n=1 is the frequency of that individual word in the corpus, for n=2 is the frequency of the bigram (sequence of 2 words) ending in, and including, that word, for n=3 is the frequency of the trigram (sequence of 3 words) ending in, and including, that word, etc. Sequences that were not found in the corpus were assigned a value of 0.

### Estimation of temporal receptive window size per electrode

We used a simplified model to simulate neural responses in the sentence (S) condition as a convolution of a stimulus train and truncated gaussian kernels with varying widths. The kernels represented an evoked ‘response function’ with a width (σ) corresponding to the temporal receptive window (TRW) of an idealized neural population underlying the intracranial responses measured by a single electrode. The kernels were constructed from gaussian curves with a standard deviation of σ/2 truncated at +/− 1 standard deviation (capturing 2/3 of the area under the gaussian). We then normalized the truncated gaussians to have a minimum of 0 and maximum of 1. We chose a symmetric kernel because we wanted to capture the full assumed TRW for a straightforward interpretation of the fitted window size. For instance, a long-tailed response functions would have a shorter “effective” receptive window because the tails of the kernel would affect the neural response much less than the center of the kernel. We further chose a kernel with smooth edges because we assumed that neural activity in response to a stimulus would increase and decrease gradually (cf. an abrupt change of voltage such as in a boxcar shape), given that macroelectrodes sum activity from a large neural population. Furthermore, note that we assumed a fixed TRW, but see <u>Discussion</u>.

We also verified that the specific shape of kernel did not affect our main result. We tested five different response functions: cosine, “wide” Gaussian (Gaussian curves with a standard deviation of σ/2 that were truncated at +/− 1 standard deviation, as used in the manuscript), “narrow” Gaussian (Gaussian curves with a standard deviation of σ/16 that were truncated at +/− 8 standard deviations), a square (i.e., boxcar) function (1 for the entire window) and a linear asymmetric function (linear function with a value of 0 initially and a value of 1 at the end of the window).

The stimulus train took a value of 1 at the time of word onsets and 0 otherwise, assuming, for simplicity, that the minimal stimulus unit of interest for language-responsive neural populations is a word (cf. e.g., Bozic et al., 2010 and Regev et al., 2021 for evidence that the language network is sensitive to sub-lexical structure). Neural responses were simulated for σ ranging from one third of a word to 8 words (the length of our stimuli), in 1 sample increments (1/27^th^ of a word, the highest resolution we were able to evaluate given our sampling rate of 60 Hz). Our implementation of the convolution is identical to assuming that the kernels appear as evoked responses starting at each word onset (see OSF https://osf.io/xfbr8/). The length of the evoked response/kernel is directly mapped onto a longer temporal receptive window, such that when a stimulus evokes a wider response its effect will remain for a longer period of time.

To find the best fit of the receptive window size for each electrode after simulating neural signals using this toy model, we selected the TRW size that yielded the highest correlation between the simulated neural response (also normalized to be between 0 and 1) and the actual neural response. The value of the correlation was taken as a proxy for the goodness of fit.

To evaluate significance, we ran linear mixed-effects (LME) models regressing the estimates temporal receptive window sizes (σ) of all electrodes on the fixed effects categorical variable of *cluster* grouped by the random effects variable of *participant*. Cluster was dummy-coded as a categorical variable with three levels, and Cluster 1 was treated as the baseline intercept. This approach allowed us to compare electrodes in Cluster 2 to those in Cluster 1, and electrodes in Cluster 3 to those in Cluster 1. To additionally compare electrodes in Clusters 2 vs. 3, we compared their LME coefficients using the Matlab procedure *coefTest*.

### Anatomical topography analysis

We examined the anatomical topographic distribution of the electrodes that exhibit the three temporal response profiles discovered in Dataset 1. Specifically, we probed the spatial relationships between all electrodes that belong to different clusters (e.g., electrodes in Cluster 1 vs. 2) with respect to the two axes: anterior-posterior [y], and inferior-superior [z]. This approach allowed us to ask whether, for example, electrodes that belong to one cluster tend to consistently fall posterior to the electrodes that belong to another cluster.

To do this, we extracted the MNI coordinates of all the electrodes in each of the three clusters and ran linear mixed-effects (LME) models regressing each of the coordinates (either y or z) on the fixed effects categorical variable of *cluster* grouped by the random effects variable of *participant*, using the Wilkinson formula:

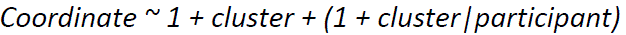

while *Coordinate* is either the y or z MNI coordinate. The random effect that groups the estimates by participant ensures that electrode coordinates are compared *within participants*. This approach is crucial for accommodating inter-individual variability in the precise locations of language areas (e.g., Fedorenko et al., 2010), which means that the absolute values of MNI coordinates cannot be easily compared between participants.

Cluster was dummy-coded as a categorical variable with three levels, and Cluster 1 was treated as the baseline intercept. This approach allowed us to compare electrodes in Cluster 2 to those in Cluster 1, and electrodes in Cluster 3 to those in Cluster 1. To additionally compare electrodes in Clusters 2 vs. 3, we ran another similar LME model with the only difference being that the baseline intercept was now the Cluster 2 category (**Tables S2-4**). To account for the small number of participants in Dataset 1, we used the Satterthwaite corrective degree-of-freedom approximation method, combined with REML fitting for LME, which was shown to be most effective when using the Satterthwaite method (Luke, 2017).

We repeated this analysis for Dataset 2, but we only examined Clusters 1 and 3, which were robustly present in that dataset. We performed the analysis for the electrodes in the two hemispheres separately.

### Replication of the clusters in Dataset 2

As described in <u>Experiment</u>, the design that was used for participants in Dataset 1 included four conditions: sentences (S), word lists (W), Jabberwocky sentences (J), and nonword lists (N). Because the design in Dataset 2 included only two of the four conditions (sentences (S) and nonword lists (N)), we first repeated the clustering procedure for Dataset 1 using only the S and N conditions to test whether similar clusters could be recovered with only a subset of conditions.

We then applied the same clustering procedure to Dataset 2 (n=16 participants, m=362 language-responsive electrodes). The elbow method revealed that the optimal number of clusters in Dataset 2 is k=2. However, because the optimal number of clusters in Dataset 1 was k=3, we examined the clustering solutions at both k=2 and k=3 levels. We also performed an analysis where we assigned electrodes in Dataset 2 to the most correlated Dataset 1 cluster. This analysis was intended to examine whether responses like those found in Dataset 1 were at all present in Dataset 2 (even if they did not emerge as strongly through clustering), and thus the assignment of electrodes to a ‘cluster’ was done by correlation alone – no actual clustering was performed.

To statistically compare the clustering solutions between Datasets 1 and 2 for k=3 and following the assignment by correlation procedure, we used the same approach as the one described above (<u>Stability of clusters across trials</u>). In particular, using Dataset 2, we shuffled average responses across electrodes (within each condition separately), re-clustered or re-assigned the electrodes into 3 clusters, and then correlated the resulting cluster averages to the cluster averages from Dataset 1. We repeated this process 1,000 times to construct a null distribution of the correlations for each of the 3 clusters. These distributions were used to calculate the probability that the correlation between the clusters across the two datasets using the actual, non-permuted Dataset 2 was higher than would be expected by chance.

To statistically compare the clustering solutions when k=3 in Dataset 1 and k=2 in Dataset 2, we used a similar procedure as the one described above. However, we only compared the resulting cluster centers from the permuted data to the two clusters in Dataset 1 that were most strongly correlated with the two clusters that emerged from Dataset 2 (i.e., Clusters 1 and 3).

## Data Availability

Preprocessed data will be publicly available on OpenNeuro at the time of publication. All stimuli and statistical results, as well as all additional analyses, are available on OSF at https://osf.io/xfbr8/. Raw data will be made available upon request.

## Code Availability

Code used to conduct analyses and generate figures from the preprocessed data will be publicly available on GitHub at the time of publication.

## Acknowledgements

We would like to acknowledge the participants for agreeing to take part in our study, as well as Nancy Kanwisher, former and current EvLab members, especially Cory Shain and Anya Ivanova, and the audience at the Neurobiology of Language conference (2022, Philadelphia) for helpful discussions and comments on the analyses and manuscript. TIR was supported by the Zuckerman-CHE STEM Leadership Program and by the Poitras Center for Psychiatric Disorders Research. CC was supported by the Kempner Institute for the Study of Natural and Artificial Intelligence at Harvard University. ALR was supported by NIH award U01-NS108916. JTW was supported by NIH awards R01-MH120194 and P41-EB018783, and the American Epilepsy Society Research & Training Fellowship for clinicians. PB was supported by NIH awards R01-EB026439, U24-NS109103, U01-NS108916, U01-NS128612 and P41-EB018783, the McDonnell Center for Systems Neuroscience and Fondazione Neurone. EF was supported by NIH awards R01-DC016607, R01-DC016950, and U01-NS121471 and research funds from the McGovern Institute for Brain Research, Brain and Cognitive Sciences Department, and the Simons Center for the Social Brain.

## Author Contribution Statement

TIR and CC equally contributed to study conception and design, data analysis and interpretation of results, and manuscript writing; EH contributed to data analysis and manuscript editing; MA to data collection and analysis; ALR, JW and PB to data collection and manuscript editing; EF contributed to study conception and design, supervision, interpretation of results, and manuscript writing.

## Supplementary Information

**Figure S1.**
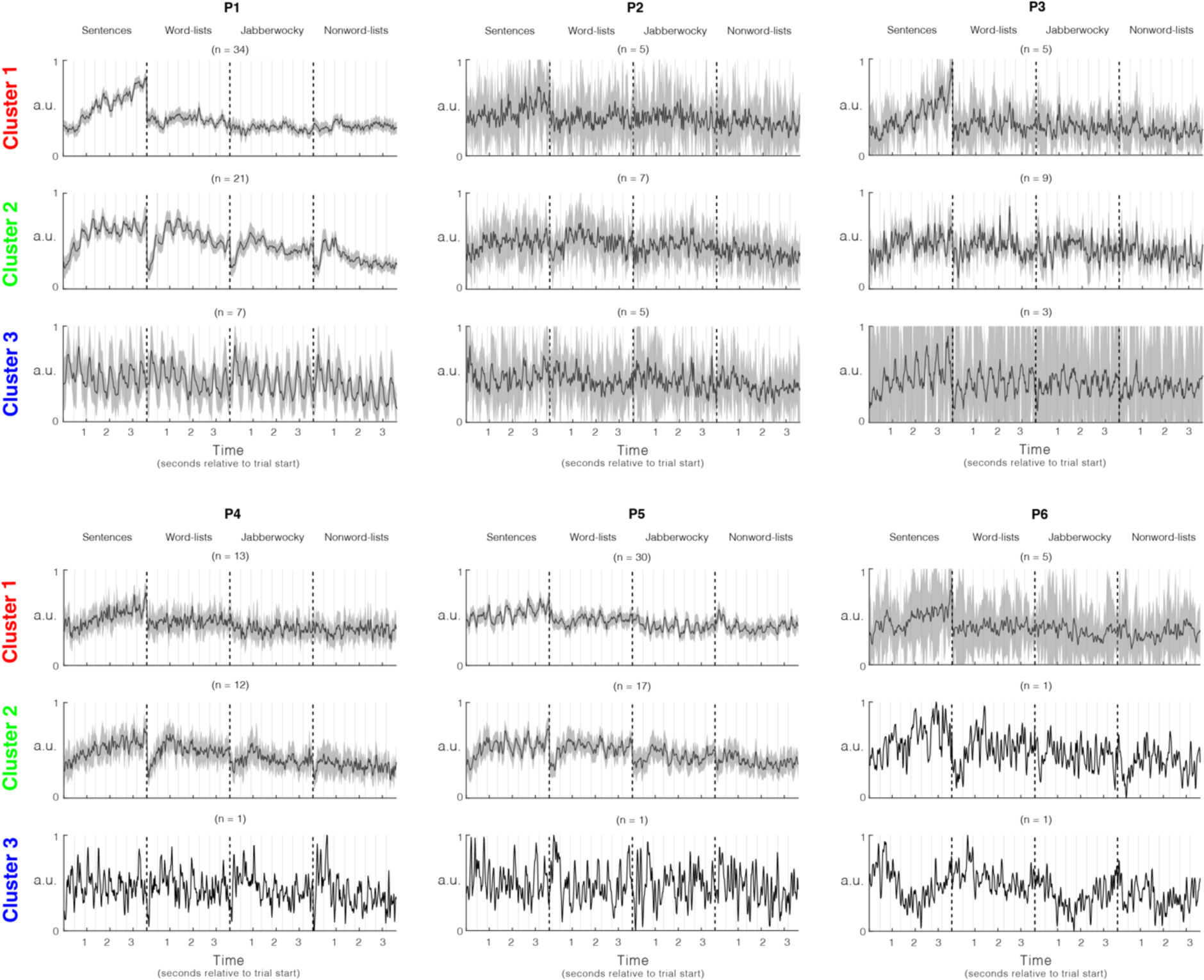
– Dataset 1 k-medoids (k=3) cluster assignments by participant. Average cluster responses as in **Figure 2E** grouped by participant. Shaded areas around the signal reflect a 99% confidence interval over electrodes. The number of electrodes constructing the average (n) is denoted above each signal in parenthesis. Prototypical responses for each of the three clusters were found in nearly all participants individually. However, for participants with only a few electrodes assigned to a given cluster (e.g., P5 Cluster 3), the responses were more variable.

**Figure S2.**
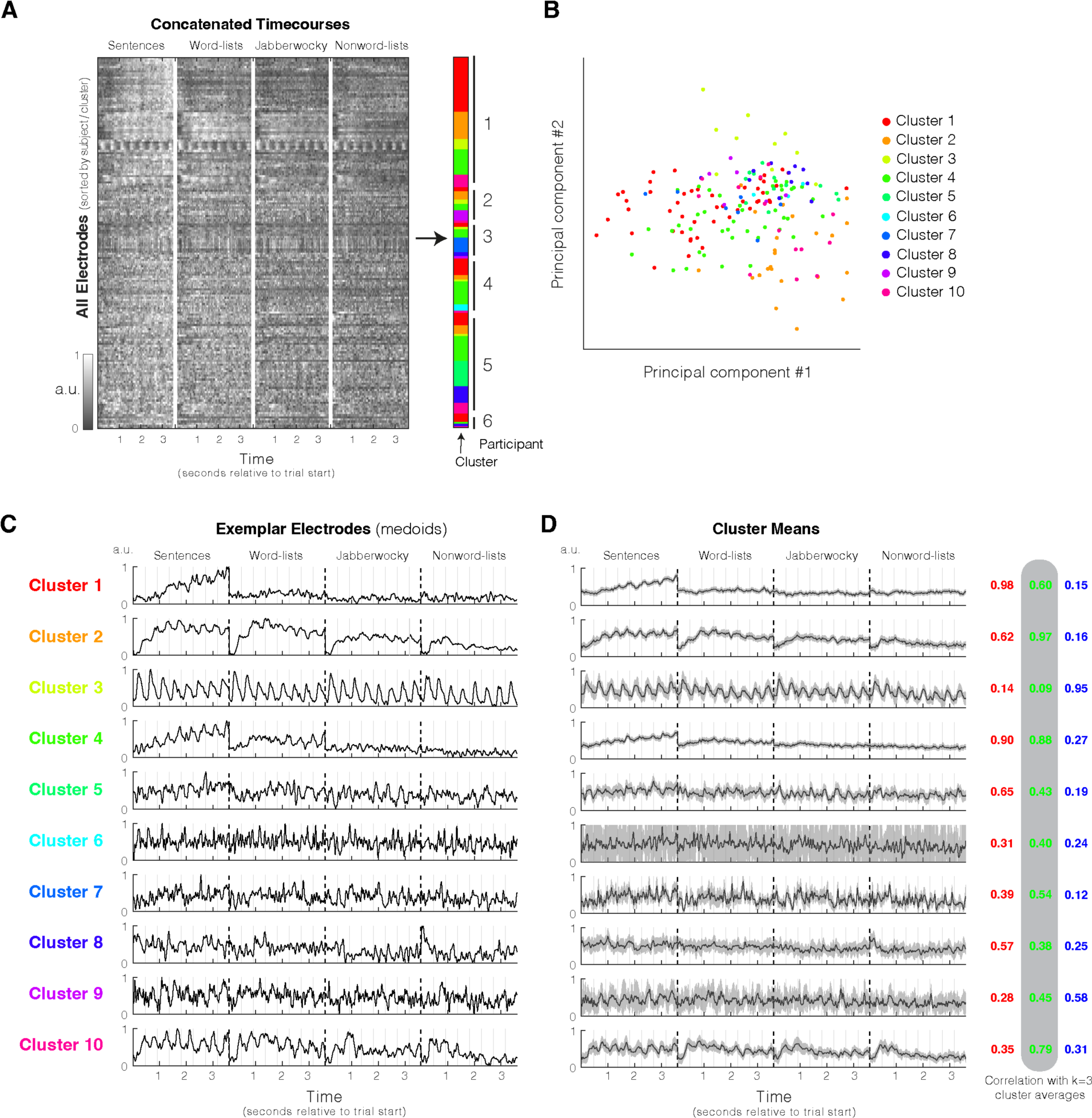
– Dataset 1 k-medoids clustering with k=10. **A**) Clustering mean electrode responses (S+W+J+N) using k-medoids (k=10) with a correlation-based distance. Shading of the data matrix reflects normalized high-gamma power (70-150Hz). **B)** Electrode responses visualized on their first two principal components, colored by cluster. **C)** Timecourses of best representative electrodes (‘medoids’) selected by the algorithm from each of the ten clusters. **D)** Timecourses averaged across all electrodes in each cluster. Shaded areas around the signal reflect a 99% confidence interval over electrodes. Correlation with the k=3 cluster averages are shown to the right of the timecourses. Many clusters exhibited high correlations with the k=3 response profiles from **Figure 2**.

**Figure S3.**
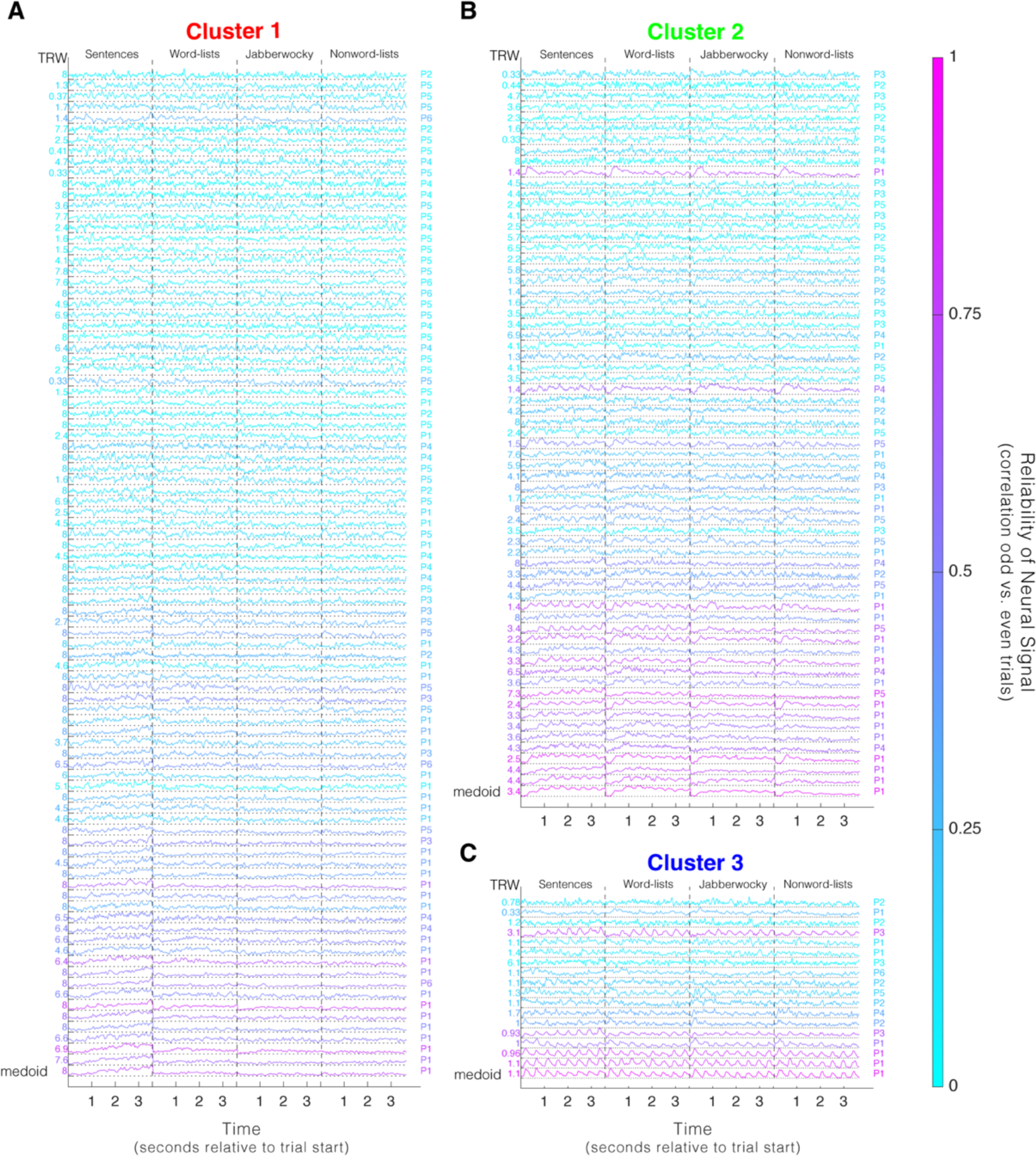
– All Dataset 1 responses. **A-C**) All Dataset 1 electrode responses. The timecourses (concatenated across the four conditions, ordered: Sentences, Word-lists, Jabberwocky, Nonword-lists) of all electrodes in Dataset 1 sorted by their correlation to the cluster medoid (shown at the bottom of each cluster). Colors reflect the reliability of the measured neural signal, computed by correlating responses to odd and even trials (**Figure 1D**). The estimated temporal receptive window (TRW) using the toy model from **Figure 4** is displayed to the left, and the participant who contributed the electrode is displayed to the right. There was strong consistency in the responses from individual electrodes within a cluster (with more variability in the less reliable electrodes), and electrodes with responses that were more similar to the cluster medoid tended to be more reliable (more pink). Note that there were two reliable response profiles (relatively pink) that showed a pattern that was distinct from the three prototypical response profiles: One electrode in Cluster 2 responded only to the onset of the first word/nonword in each trial; and one electrode in Cluster 3 was highly locked to all onsets *except* the first word/nonword. These profiles indicate that although the prototypical clusters explain a substantial amount of the functional heterogeneity of responses in the language network, they were not the *only* observed responses.

**Figure S4.**
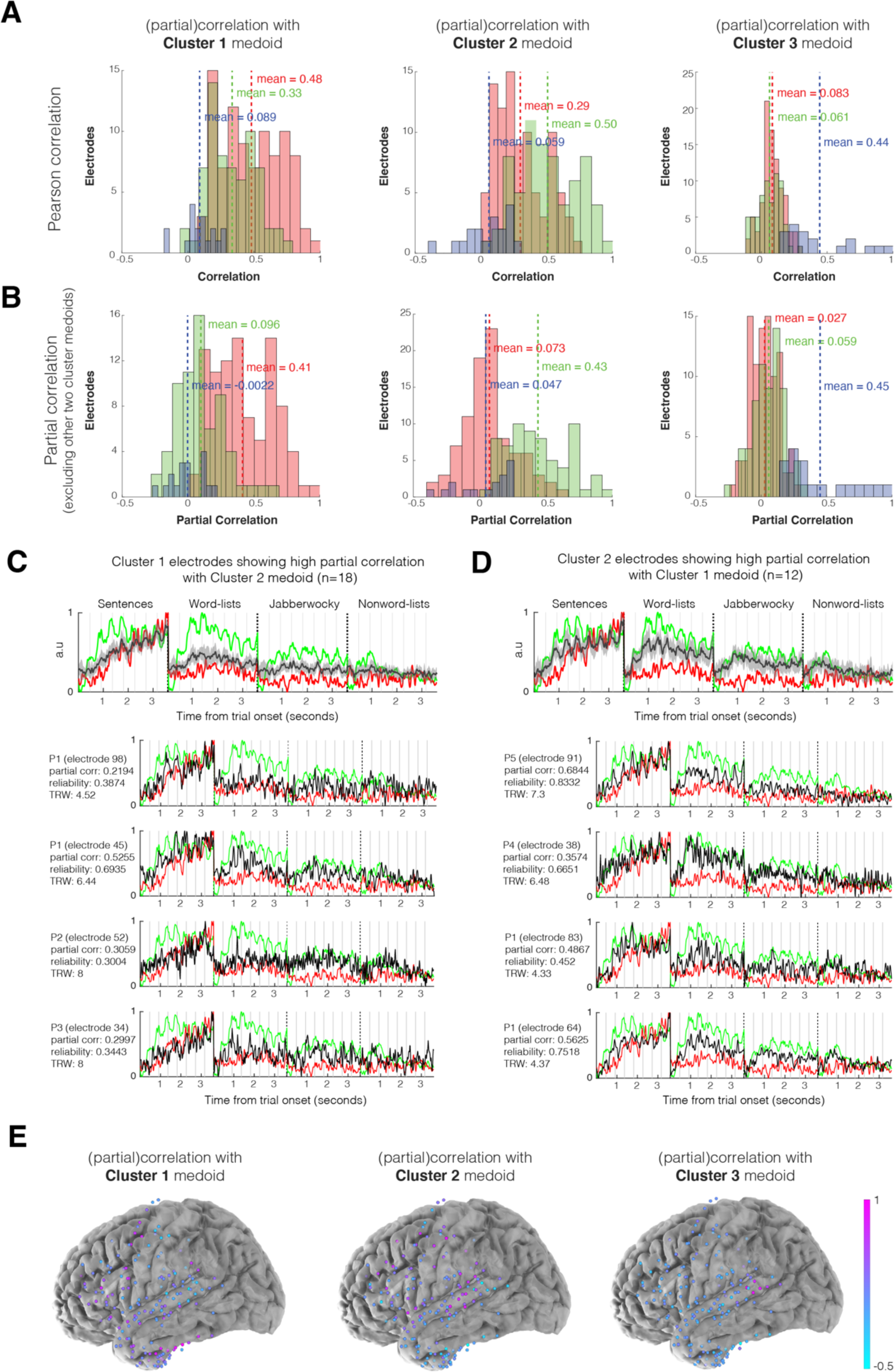
– Partial correlations of individual response profiles with each of the cluster medoids. **A**) Pearson correlations of all response profiles with each of the cluster medoids, grouped by cluster assignment. **B)** Partial correlations (<u>Methods</u>) of all response profiles with each of the cluster medoids, controlling for the other two cluster medoids, grouped by cluster assignment. **C)** Response profiles from electrodes assigned to Cluster 1 that had a high partial correlation (>0.2, arbitrarily defined) with the Cluster 2 medoid (and split-half reliability>0.3). ***Top:*** Average over all electrodes that met these criteria (n=18, black). The Cluster 1 medoid is shown in red, and the Cluster 2 medoid is shown in green. ***Bottom:*** Four sample electrodes (black). **D)** Response profiles assigned to Cluster 2 that had a high partial correlation (>0.2, arbitrarily defined) with the Cluster 1 medoid (and split-half reliability>0.3). ***Top*:** Average over all electrodes that meet these criteria (n=12, black). The Cluster 1 medoid is shown in red, and the Cluster 2 medoid is shown in green. ***Bottom:*** Four sample electrodes (black; see osf.io/xfbr8/ for all mixed response profiles with split-half reliability>0.3). **E)** Anatomical distribution of electrodes in Dataset 1 colored by their partial correlation with a given cluster medoid (controlling for the other two medoids). Cluster-1– and Cluster-2-like responses were present throughout frontal and temporal areas (with Cluster 1 responses having a slightly higher concentration in the temporal pole and Cluster 2 responses having a slightly higher concentration in the superior temporal gyrus (STG)), whereas Cluster-3-like responses were localized to the posterior STG.

**Figure S5.**
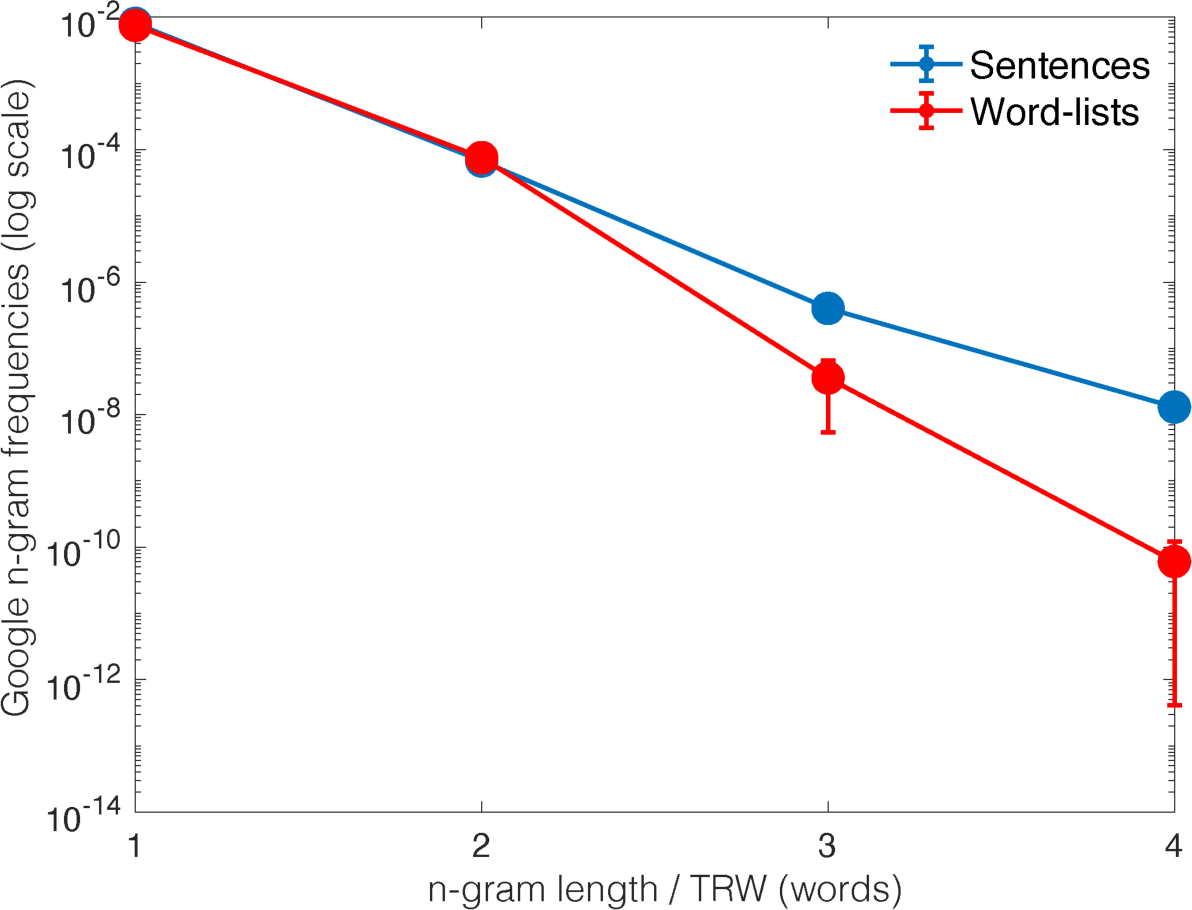
– N-gram frequencies of sentences and word lists diverge with n-gram length. N-gram frequencies were extracted from the Google n-gram online platform (https://books.google.com/ngrams/), averaging across Google books corpora between the years 2010 and 2020. For each individual word, the n-gram frequency for n=1 was the frequency of that word in the corpus; for n=2 it was the frequency of the bigram (sequence of 2 words) ending in that word; for n=3 it was the frequency of the trigram (sequence of 3 words) ending in that word; an so on. Sequences that were not found in the corpus were assigned a value of 0. Results are only presented until n=4 because for n>4 most of the string sequences, both from the Sentence and Word-list conditions, were not found in the corpora. The plot shows that the difference between the log n-gram values for the sentences and wordlists in our stimulus set grew as a function of N.

**Figure S6.**
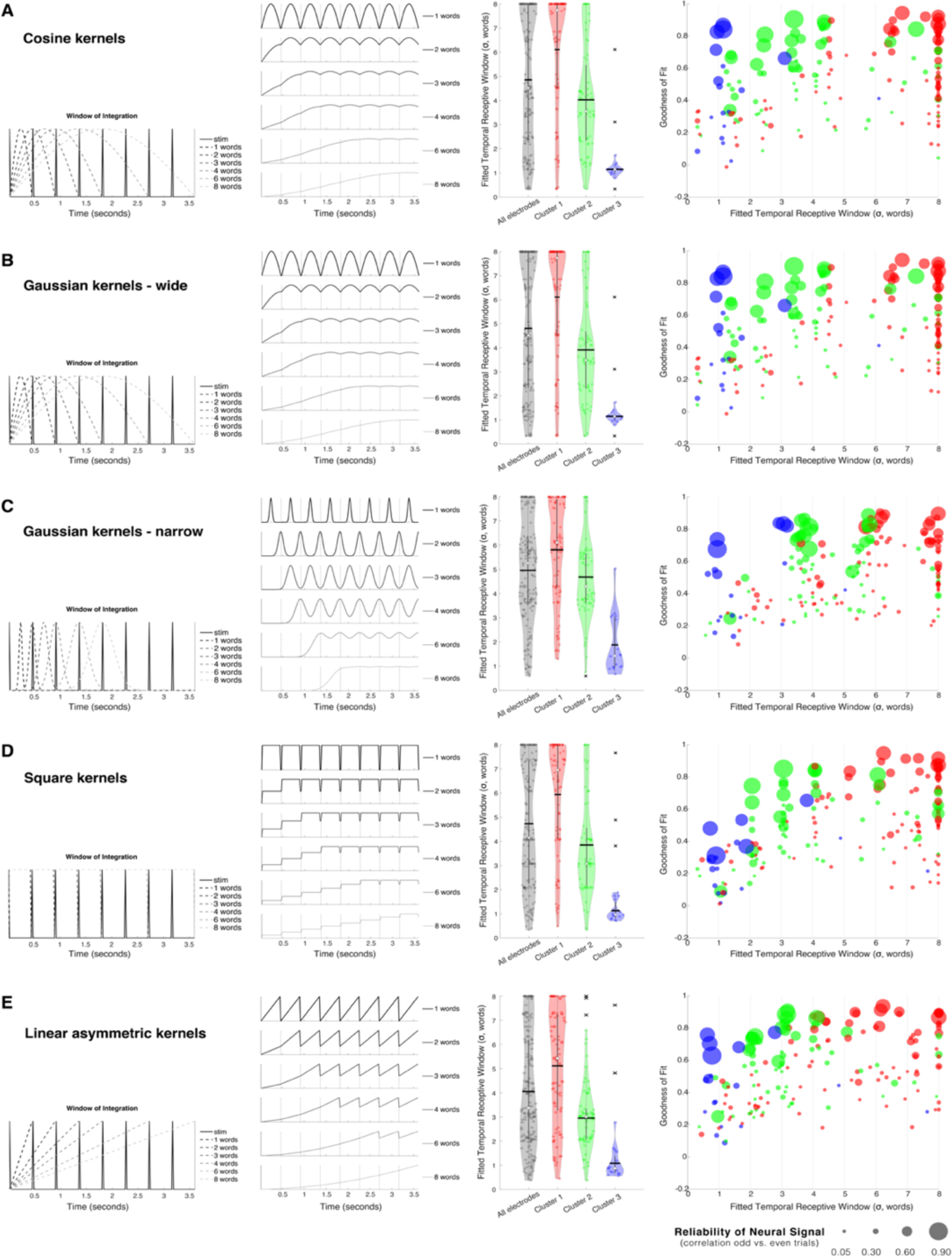
– Temporal receptive window (TRW) estimates with kernels of different shapes. The toy TRW model from **Figure 4** was applied using five different kernel shapes: cosine (**A**), “wide” Gaussian (Gaussian curves with a standard deviation of σ/2 that were truncated at +/− 1 standard deviation, as used in **Figure 4**; **B**), “narrow” Gaussian (Gaussian curves with a standard deviation of σ/16 that were truncated at +/− 8 standard deviations; **C**), a square (i.e., boxcar) function (1 for the entire window; **D**) and a linear asymmetric function (linear function with a value of 0 initially and a value of 1 at the end of the window; **E**). For each kernel (**A-E**), the plots represent (left to right, all details are identical to **Figure 4** in the manuscript): 1) The kernel shapes for TRW = 1, 2, 3, 4, 6 and 8 words, superimposed on the simplified stimulus train; 2) The simulated neural signals for each of those TRWs; 3) violin plots of best fitted TRW values across electrodes (each dot represents and electrode) for all electrodes (black), or electrodes from only Clusters 1 (red) 2 (green) or 3 (blue); and 4) Estimated TRW as a function of goodness of fit. Each dot is an electrode, its size represents the reliability of its neural response, computed via correlation between the mean signals when using only odd vs. only even trials, x-axis is the electrode’s best fitted TRW, y-axis is the goodness of fit, computed via correlation between the neural signal and the closest simulated signal. For all kernels the TRWs showed a decreasing trend from Cluster 1 to 3.

**Figure S7.**
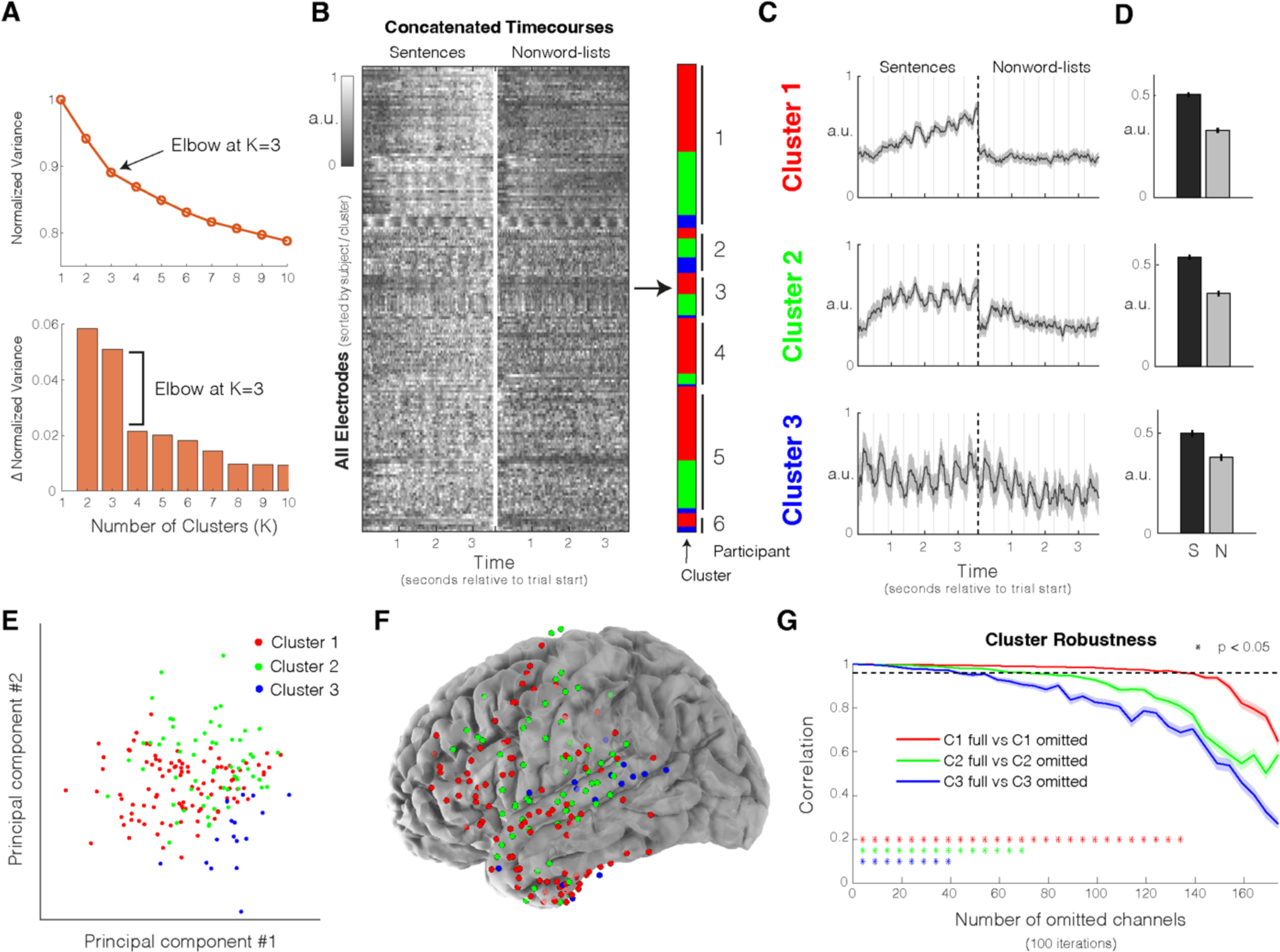
– Dataset 1 k-medoids clustering results with only S-N conditions. **A**) Search for optimal k using the “elbow method”. ***Top:*** variance (sum of the distances of all electrodes to their assigned cluster center) normalized by the variance when k=1 as a function of k (normalized variance (NV)). ***Bottom:*** change in NV as a function of k (NV(k+1) – NV(k)). After k=3 the change in variance became more moderate, suggesting that 3 clusters appropriately described Dataset 1 when using only the responses to sentences and nonwords (as was the case when all four conditions were used). **B)** Clustering mean electrode responses (only S+N, importantly) using k-medoids (k=3) with a correlation-based distance. Shading of the data matrix reflects normalized high-gamma power (70-150Hz). **C)** Average timecourse by cluster. Shaded areas around the signal reflect a 99% confidence interval over electrodes. Clusters 1-3 showed a strong similarity to the clusters reported in **Figure 2**. **D)** Mean condition responses by cluster. Error bars reflect standard error of the mean over electrodes. **E)** Electrode responses visualized on their first two principal components, colored by cluster. **F)** Anatomical distribution of clusters across all participants (n=6). **G)** Robustness of clusters to electrode omission (random subsets of electrodes were removed in increments of 5). Stars reflect significant similarity with the full dataset (p<0.05; evaluated with a permutation test; <u>Methods</u>). Shaded regions reflect standard error of the mean over randomly sampled subsets of electrodes. Relative to when all conditions were used, Cluster 2 was less robust to electrode omission (although still more robust than Cluster 3), suggesting that responses to word lists and Jabberwocky sentences (both not present here) are particularly important for distinguishing Cluster 2 electrodes from Cluster 1 and 3 electrodes.

**Figure S8.**
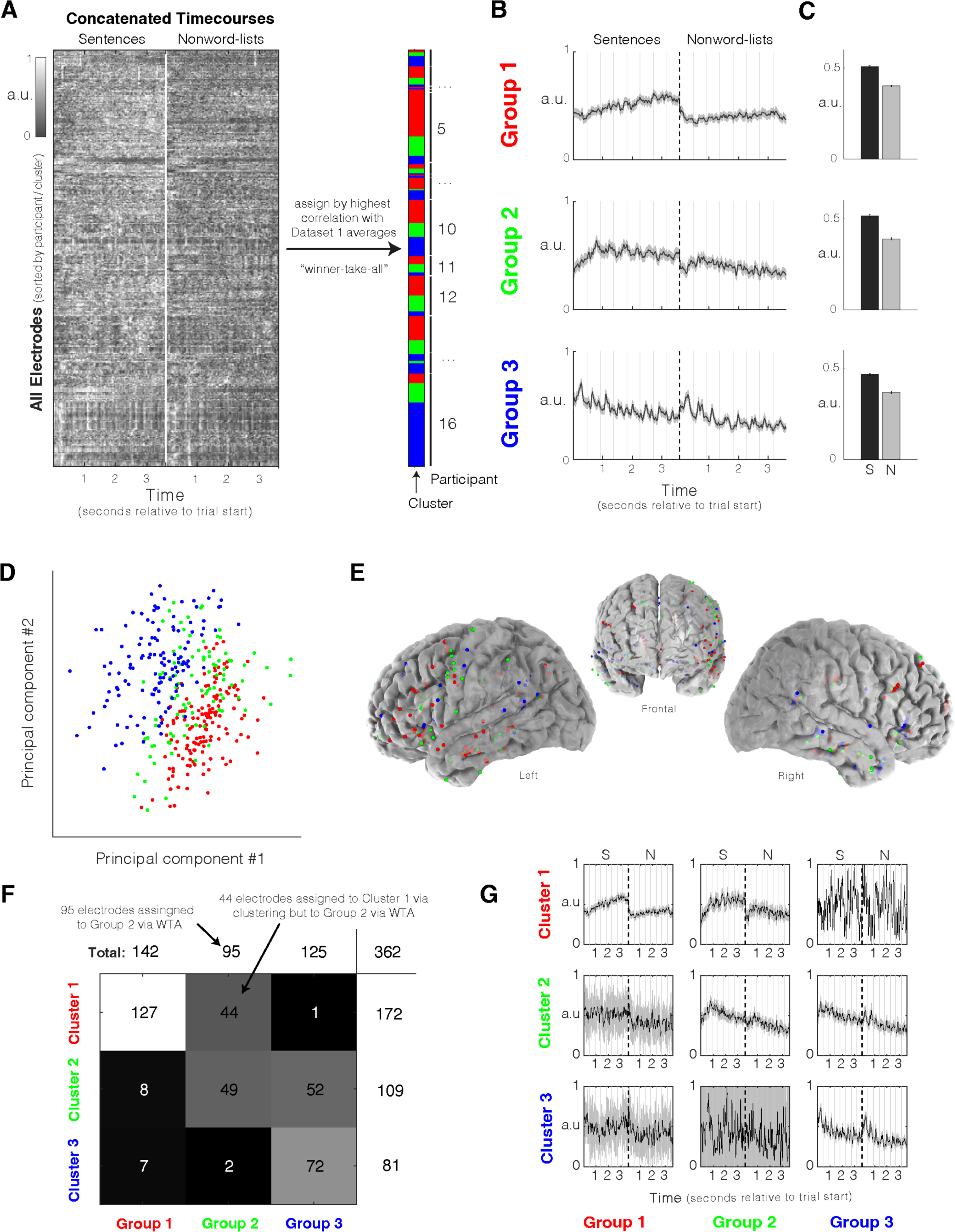
– Dataset 2 electrode assignment to most correlated Dataset 1 cluster under “winner– take-all” (WTA) approach. **A**) Assigning electrodes from Dataset 2 to the most correlated cluster from Dataset 1. Assignment was performed using the correlation with the Dataset 1 cluster average, not the cluster medoid. Shading of the data matrix reflects normalized high-gamma power (70-150Hz). **B)** Average timecourse by group. Shaded areas around the signal reflect a 99% confidence interval over electrodes. **C)** Mean condition responses by group. Error bars reflect standard error of the mean over electrodes. **D)** Electrode responses visualized on their first two principal components, colored by group. **E)** Anatomical distribution of groups across all participants (n=16). **F-G)** Comparison of cluster assignment of electrodes from Dataset 2 using clustering vs. winner-take-all (WTA) approach. **F)** The numbers in the matrix correspond to the number of electrodes assigned to cluster y during clustering (y-axis) versus the number electrodes assigned to group x during the WTA approach (x-axis). For instance, there were 44 electrodes that were assigned to Cluster 1 during clustering but were “pulled out” to Group 2 (the analog of Cluster 2) during the WTA approach. The total number of electrodes assigned to each cluster during the clustering approach are shown to the right of each row. The total number of electrodes assigned to each group during the WTA approach are shown at the top of each column. N=362 is the total number of electrodes in Dataset 2. **G)** Similar to **F**, but here the average timecourse across all electrodes assigned to the same cluster/group during both procedures is presented. Shaded areas around the signals reflect a 99% confidence interval over electrodes.

**Figure S9.**
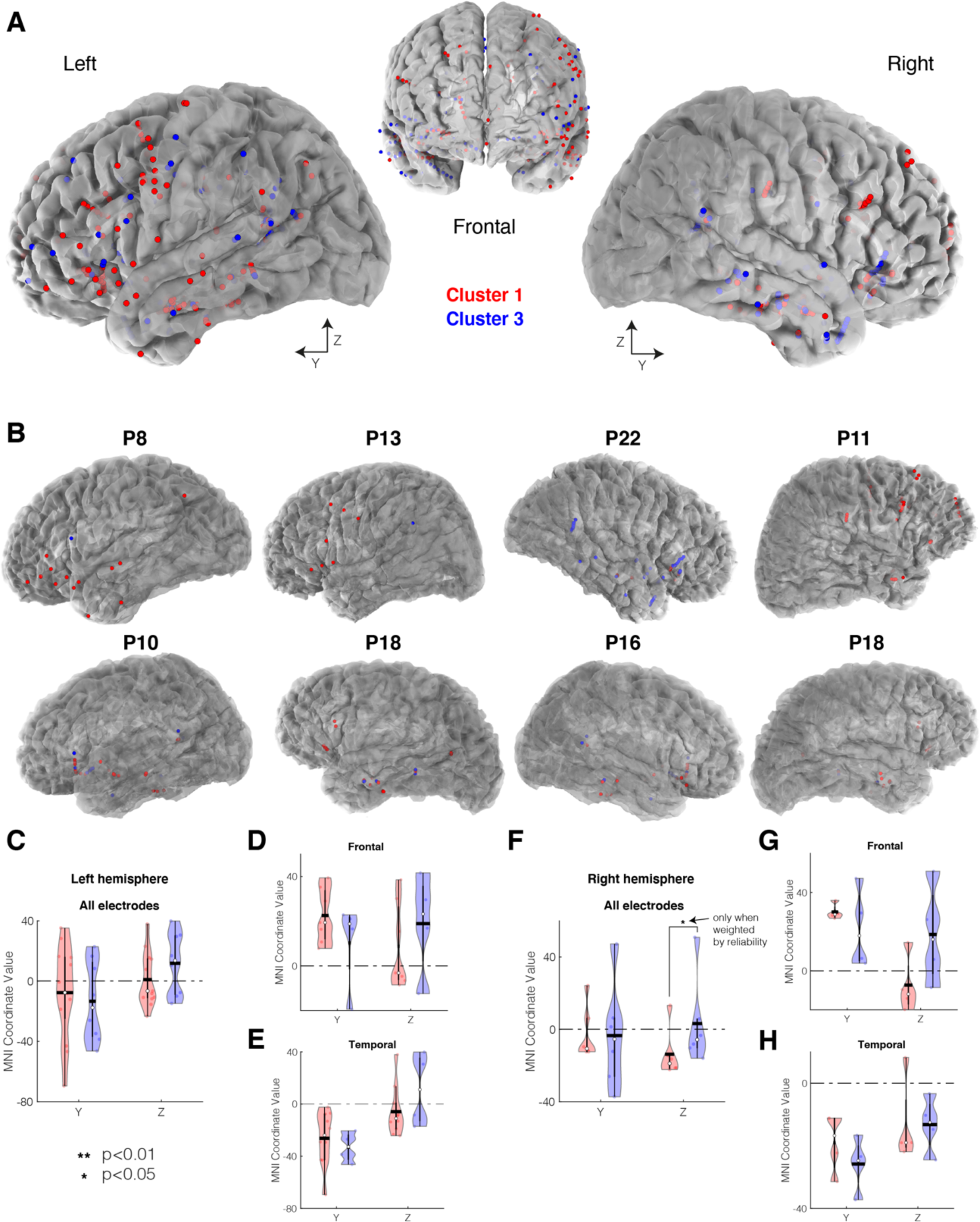
– Anatomical distribution of the clusters in Dataset 2. Anatomical distribution of language-responsive electrodes in Dataset 2 across all subjects in MNI space, colored by cluster. Only Clusters 1 and 3 (those from Dataset 1 that replicate to Dataset 2) are shown. **B)** Anatomical distribution of language-responsive electrodes in subject-specific space for eight sample participants. **C-H)** Violin plots of MNI coordinate values for Clusters 1 and 3 in the left and right hemisphere (**C-E** and **F-H**, respectively), where plotted points represent the mean of all coordinate values for a given participant and cluster. The mean is plotted with a black horizontal line, and the median is shown with a white circle. Significance is evaluated with a LME model (<u>Methods</u>, **Tables S3** and **S4**). The Cluster 3 posterior bias from Dataset 1 was weakly present but not statistically reliable.

**Figure S10.**
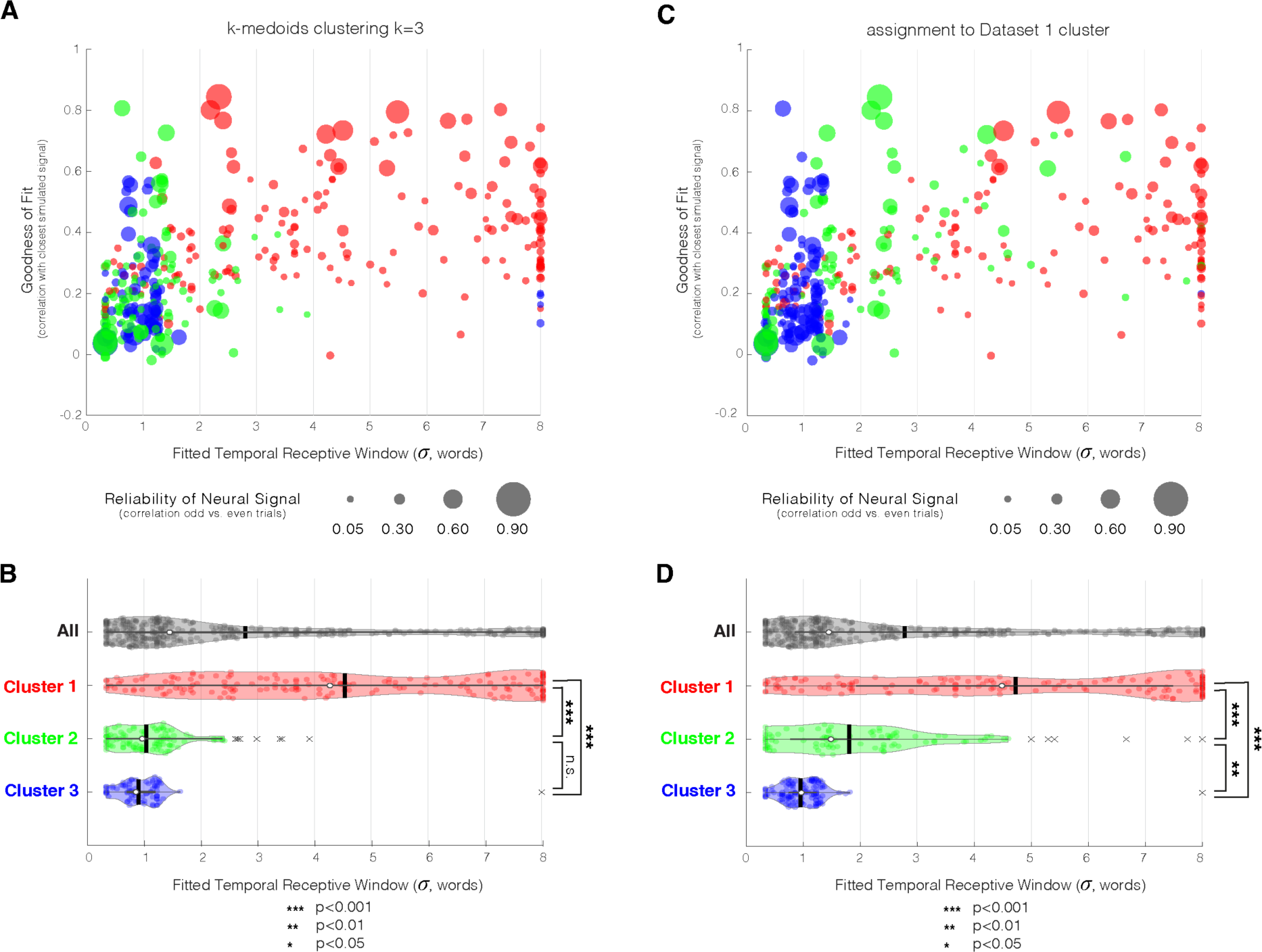
– Estimation of temporal receptive window (TRW) sizes for electrodes in Dataset 2. As in **Figure 4** but for electrodes in Dataset 2. **A)** Best TRW fit (using the toy model from **Figure 4**) for all electrodes, colored by cluster (when k-medoids clustering with k=3 was applied, **Figure 6**) and sized by the reliability of the neural signal as estimated by correlating responses to odd and even trials (**Figure 6C**). The ‘goodness of fit’, or correlation between the simulated and observed neural signal (Sentence condition only), is shown on the y-axis. **B)** Estimated TRW sizes across all electrodes (grey) and per cluster (red, green, and blue). Black vertical lines correspond to the mean window size and the white dots correspond to the median. “x” marks indicate outliers (more than 1.5 interquartile ranges above the upper quartile or less than 1.5 interquartile ranges below the lower quartile). Significance values were calculated using a linear mixed-effects model (<u>Methods</u>, **Table S8**). **C-D)** Same as **A** and **B**, respectively, except clusters were assigned by highest correlation with Dataset 1 clusters (**Figure S8**). Under this procedure, Cluster 2 reliably separated from Cluster 3 in terms of its TRW (all ps<0.001, evaluated with a LME model, <u>Methods</u>, **Table S9**).

**Table S1A.** LME results quantifying degree of stimulus locking by cluster. All estimates from the linear mixed-effects model (LME) regressing the locking value (<u>Methods</u>) on the categorical variables of cluster (3 levels) and condition (4 levels for Sentences (S), Word-lists (W), Jabberwocky (J), Nonword-lists (N), <u>Methods</u>), including their interaction, all grouped by the random variable of participant. Model formula: *Locking ∼ cluster*condition + (cluster|participant) + (condition|participant)*). The Satterthwaite Method was used to estimate the degrees of freedom (DF) due to our small sample size. Implemented with Matlab *fitlme* routine. Semicolon represents interactions. The intercept represents one level of each of the categorical variables and is denoted by “reference”. The models are reference-coded such that all estimate values are evaluated and compared statistically to the intercept/reference. Two models are presented, separated by a horizontal line. The only difference between the models regards the level of the categorical variable ‘Cluster’ that is assigned to the reference. In the first model the intercept/reference is for Cluster 1 (C1), and in the second model it is Cluster 2 (C2). We used the second model in order to obtain the statistical comparisons between clusters 2 and 3. The estimate magnitudes show a trend for stimulus locking by cluster: C1<C2<C3, but the only pairwise comparison that reached significance was of Cluster 3 to 1 (p<0.05) and the other comparisons were marginally significant (p<0.1). Estimate magnitudes further show a trend for stimulus locking by Condition: S>W>J>N, but the only pairwise comparison that reached significance was S vs. N (p<0.05) in the first model, and J vs. S in the second model (p<0.001). No interaction terms were significant. An additional ANOVA test for LME revealed a significant main effect for cluster (F(2,9.13)=5.4, p<0.05) and the main effect for condition as well as the interaction term did not reach significance. See **Figure 5**.

| Name | Estimate | SE | tStat | DF | pValue |
| --- | --- | --- | --- | --- | --- |
| **C1, Condition S – reference** | **0.103** | **0.023** | **4.503** | **10.332** | **0.001** |
| C2 – relative to C1 | 0.074 | 0.038 | 1.974 | 11.573 | 0.073 |
| **C3 – relative to C1** | **0.239** | **0.080** | **2.971** | **7.464** | **0.019** |
| Condition W – relative to S | -0.023 | 0.026 | -0.880 | 681.988 | 0.38 |
| Condition J – relative to S | -0.036 | 0.027 | -1.338 | 27.077 | 0.19 |
| **Condition N – relative to S** | **-0.060** | **0.027** | **-2.270** | **46.806** | **0.028** |
| Condition W: C2 relative to C1 | -0.026 | 0.040 | -0.657 | 682.349 | 0.51 |
| Condition W: C3 relative to C1 | 0.022 | 0.063 | 0.348 | 682.352 | 0.73 |
| Condition J: C2 relative to C1 | -0.041 | 0.040 | -1.046 | 677.506 | 0.30 |
| Condition J: C3 relative to C1 | 0.007 | 0.064 | 0.111 | 644.768 | 0.91 |
| Condition N: C2 relative to C1 | -0.063 | 0.040 | -1.588 | 680.554 | 0.11 |
| Condition N: C3 relative to C1 | -0.046 | 0.063 | -0.718 | 665.329 | 0.47 |
| **C2, Condition S – reference** | **0.178** | **0.029** | **6.053** | **11.092** | **0.00008** |
| C3 – relative to C2 | 0.164 | 0.082 | 2.001 | 7.518 | 0.083 |
| C1 – relative to C2 | -0.074 | 0.038 | -1.974 | 11.573 | 0.073 |
| Condition W – relative to S | -0.049 | 0.030 | -1.614 | 682.210 | 0.11 |
| **Condition J – relative to S** | **-0.078** | **0.031** | **-2.485** | **52.580** | **0.016** |
| **Condition N – relative to S** | **-0.123** | **0.031** | **-4.000** | **90.665** | **0.00013** |
| Condition W: C3 relative to C2 | 0.048 | 0.065 | 0.735 | 682.350 | 0.46 |
| Condition W: C1 relative to C2 | 0.026 | 0.040 | 0.657 | 682.349 | 0.51 |
| Condition J: C3 relative to C2 | 0.048 | 0.065 | 0.741 | 669.544 | 0.46 |
| Condition J: C1 relative to C2 | 0.041 | 0.040 | 1.046 | 677.506 | 0.30 |
| Condition N: C3 relative to C2 | 0.017 | 0.065 | 0.262 | 677.099 | 0.79 |
| Condition N: C1 relative to C2 | 0.063 | 0.040 | 1.588 | 680.554 | 0.11 |

**Table S1B.** ANOVA for the LME results presented in Table S1A. ANOVA for LME was run on the first model presented in Table S1A. These results reveal that the main effect of Cluster was overall significant (p<0.05), but the main effect of Condition as well as the interaction between cluster and condition did not reach significance.

| Name | FStat | DF1 | DF2 | pValue |
| --- | --- | --- | --- | --- |
| **(Intercept)** | **20.3** | **1** | **10.33** | **0.0010** |
| **Cluster** | **5.4** | **2** | **9.13** | **0.028** |
| Condition | 1.9 | 3 | 41.84 | 0.15 |
| Cluster : Condition | 0.6 | 6 | 670.00 | 0.73 |

**Table S2A.**
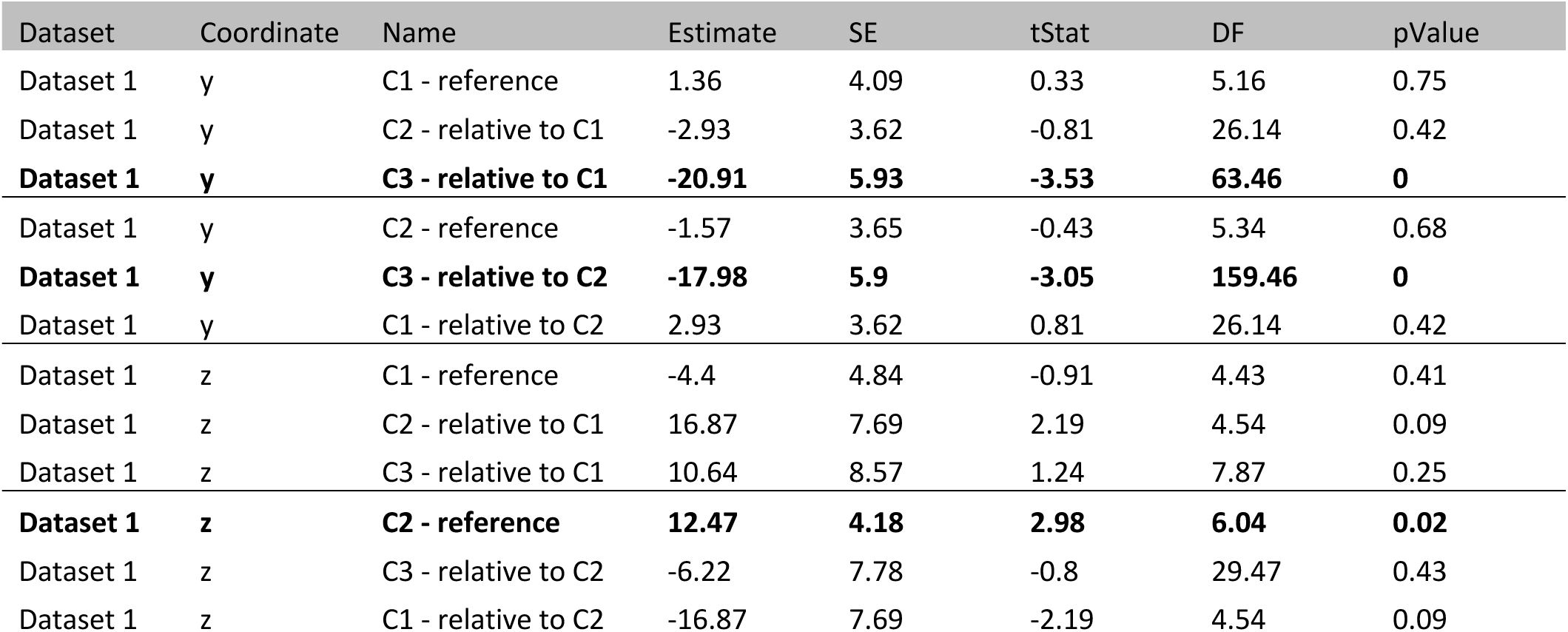
– LME results comparing MNI coordinates of the 3 clusters, Dataset 1, Left hemisphere. All estimates from the linear mixed-effects model (LME) regressing the y (posterior-anterior) and z (inferior-superior) MNI coordinates (<u>Methods</u>) on the categorical variable of cluster (3 levels) grouped by the random variable of participant. Model formula: *MNI coordinate ∼ cluster + (cluster|participant).* Details are similar to Table S1A. The y-coordinate of Cluster 3 was significantly different from Clusters 1 and 2 (ps<0.01). All the other comparisons did not reach significance. See **Figure 6**.

**Table S2B.** – Same as Table S2A but electrodes were weighted by reliability.

| Dataset | Coordinate | Name | Estimate | SE | tStat | DF | pValue |
| --- | --- | --- | --- | --- | --- | --- | --- |
| Dataset 1 | y | C1 - reference | 4.9 | 5.6 | 0.9 | 5.2 | 0.413759 |
| **Dataset 1** | **y** | **C2 - relative to C1** | **-8.3** | **3.7** | **-2.2** | **38.1** | **0.031376** |
| **Dataset 1** | **y** | **C3 - relative to C1** | **-27.9** | **5.4** | **-5.1** | **166.1** | **8.00E-07** |
| Dataset 1 | y | C2 - reference | -3.4 | 4.9 | -0.7 | 4.9 | 0.52027 |
| **Dataset 1** | **y** | **C3 - relative to C2** | **-19.5** | **5.5** | **-3.6** | **64.7** | **0.000706** |
| **Dataset 1** | **y** | **C1 - relative to C2** | **8.3** | **3.7** | **2.2** | **38.1** | **0.031376** |
| Dataset 1 | z | C1 - reference | -1.3 | 3.4 | -0.4 | 3 | 0.732998 |
| **Dataset 1** | **z** | **C2 - relative to C1** | **19.2** | **5.1** | **3.8** | **2.5** | **0.045907** |
| Dataset 1 | z | C3 - relative to C1 | 14.8 | 7.5 | 2 | 2.5 | 0.162557 |
| **Dataset 1** | **z** | **C2 - reference** | **17.9** | **3.9** | **4.6** | **1.7** | **0.059805** |
| Dataset 1 | z | C3 - relative to C2 | -4.4 | 8.7 | -0.5 | 1.7 | 0.672232 |
| **Dataset 1** | **z** | **C1 - relative to C2** | **-19.2** | **5.1** | **-3.8** | **2.5** | **0.045907** |

**Table S2C.** – Same as Table S2A but only frontal electrodes.

| Dataset | Coordinate | Name | Estimate | SE | tStat | DF | pValue |
| --- | --- | --- | --- | --- | --- | --- | --- |
| Dataset 1 | y | C1 - reference | 9.6 | 7.6 | 1.3 | 5.3 | 0.26 |
| Dataset 1 | y | C2 - relative to C1 | -5.6 | 6.0 | -0.9 | 5.0 | 0.40 |
| Dataset 1 | y | C3 - relative to C1 | -10.9 | 11.0 | -1.0 | 2.4 | 0.41 |
| Dataset 1 | y | C2 - reference | 4.0 | 5.0 | 0.8 | 4.7 | 0.46 |
| Dataset 1 | y | C3 - relative to C2 | -5.3 | 11.0 | -0.5 | 4.5 | 0.65 |
| Dataset 1 | y | C1 - relative to C2 | 5.6 | 6.0 | 0.9 | 5.0 | 0.40 |
| **Dataset 1** | **z** | **C1 - reference** | **13.3** | **4.7** | **2.8** | **6.7** | **0.03** |
| Dataset 1 | z | C2 - relative to C1 | 6.2 | 7.7 | 0.8 | 4.7 | 0.46 |
| Dataset 1 | z | C3 - relative to C1 | 8.5 | 11.5 | 0.7 | 10.1 | 0.48 |
| Dataset 1 | z | C2 - reference | 19.5 | 9.0 | 2.2 | 4.0 | 0.10 |
| Dataset 1 | z | C3 - relative to C2 | 2.3 | 14.5 | 0.2 | 4.5 | 0.88 |
| Dataset 1 | z | C1 - relative to C2 | -6.2 | 7.7 | -0.8 | 4.7 | 0.46 |

**Table S2D.** – Same as Table S2A but only temporal electrodes.

| Dataset | Coordinate | Name | Estimate | SE | tStat | DF | pValue |
| --- | --- | --- | --- | --- | --- | --- | --- |
| Dataset 1 | y | C1 - reference | -3.8 | 4.5 | -0.8 | 3.7 | 0.45 |
| Dataset 1 | y | C2 - relative to C1 | 0.4 | 4.4 | 0.1 | 42.8 | 0.93 |
| **Dataset 1** | **y** | **C3 - relative to C1** | **-22.3** | **6.9** | **-3.2** | **26.6** | **0.003** |
| Dataset 1 | y | C2 - reference | -3.4 | 5.6 | -0.6 | 3.8 | 0.58 |
| **Dataset 1** | **y** | **C3 - relative to C2** | **-22.7** | **7.5** | **-3.0** | **17.6** | **0.008** |
| Dataset 1 | y | C1 - relative to C2 | -0.4 | 4.4 | -0.1 | 42.8 | 0.93 |
| Dataset 1 | z | C1 - reference | -6.4 | 13.8 | -0.5 | 3.6 | 0.67 |
| Dataset 1 | z | C2 - relative to C1 | 18.2 | 7.5 | 2.4 | 1.7 | 0.16 |
| Dataset 1 | z | C3 - relative to C1 | 10.4 | 15.9 | 0.7 | 1.7 | 0.59 |
| Dataset 1 | z | C2 - reference | 11.8 | 9.5 | 1.2 | 3.1 | 0.30 |
| Dataset 1 | z | C3 - relative to C2 | -7.8 | 11.6 | -0.7 | 1.7 | 0.58 |
| Dataset 1 | z | C1 - relative to C2 | -18.2 | 7.5 | -2.4 | 1.7 | 0.16 |

**Table S3A.** – LME results comparing coordinates of the 3 clusters, Dataset 2, Left hemisphere. Similar to Table S2A but for Dataset 2, left hemisphere electrodes. The only significant comparison was the z-coordinate of Cluster 2 relative to Clusters 1 (p<0.01). See **Figure S5**.

| Dataset | Coordinate | Name | Estimate | SE | tStat | DF | pValue |
| --- | --- | --- | --- | --- | --- | --- | --- |
| Dataset 2 | y | C1 - reference | -5.3 | 7.5 | -0.7 | 8.9 | 0.5 |
| Dataset 2 | y | C2 - relative to C1 | -7.2 | 10.4 | -0.7 | 7.2 | 0.5 |
| Dataset 2 | y | C3 - relative to C1 | -1.5 | 11.0 | -0.1 | 7.8 | 0.9 |
| Dataset 2 | y | C2 - reference | -12.6 | 7.8 | -1.6 | 10.4 | 0.1 |
| Dataset 2 | y | C3 - relative to C2 | 5.7 | 8.3 | 0.7 | 9.4 | 0.5 |
| Dataset 2 | y | C1 - relative to C2 | 7.2 | 10.4 | 0.7 | 7.2 | 0.5 |
| Dataset 2 | z | C1 - reference | 3.1 | 4.2 | 0.7 | 12.6 | 0.5 |
| **Dataset 2** | **z** | **C2 - relative to C1** | **12.8** | **3.9** | **3.3** | **183.7** | **0.001** |
| Dataset 2 | z | C3 - relative to C1 | 7.3 | 5.9 | 1.2 | 30.1 | 0.2 |
| **Dataset 2** | **z** | **C2 - reference** | **15.8** | **4.6** | **3.4** | **12.0** | **0.005** |
| Dataset 2 | z | C3 - relative to C2 | -5.4 | 5.9 | -0.9 | 38.8 | 0.4 |
| **Dataset 2** | **z** | **C1 - relative to C2** | **-12.8** | **3.9** | **-3.3** | **183.7** | **0.001** |

**Table S3B.** – Same as Table S3A but electrodes are weighted by reliability.

| Dataset | Coordinate | Name | Estimate | SE | tStat | DF | pValue |
| --- | --- | --- | --- | --- | --- | --- | --- |
| Dataset 2 | y | C1 - reference | -0.1 | 6.3 | 0.0 | 7.2 | 0.98 |
| Dataset 2 | y | C2 - relative to C1 | -16.2 | 11.6 | -1.4 | 6.7 | 0.21 |
| Dataset 2 | y | C3 - relative to C1 | -4.6 | 10.4 | -0.4 | 5.1 | 0.68 |
| Dataset 2 | y | C2 - reference | -16.3 | 8.5 | -1.9 | 8.4 | 0.09 |
| Dataset 2 | y | C3 - relative to C2 | 11.6 | 8.9 | 1.3 | 9.5 | 0.23 |
| Dataset 2 | y | C1 - relative to C2 | 16.2 | 11.6 | 1.4 | 6.7 | 0.21 |
| Dataset 2 | z | C1 - reference | 4.2 | 5.0 | 0.8 | 11.7 | 0.42 |
| **Dataset 2** | **z** | **C2 - relative to C1** | **15.4** | **5.8** | **2.6** | **11.0** | **0.02** |
| Dataset 2 | z | C3 - relative to C1 | 14.0 | 7.2 | 1.9 | 8.2 | 0.09 |
| **Dataset 2** | **z** | **C2 - reference** | **19.5** | **4.7** | **4.2** | **9.3** | **0.002** |
| Dataset 2 | z | C3 - relative to C2 | -1.4 | 5.9 | -0.2 | 11.4 | 0.82 |
| **Dataset 2** | **z** | **C1 - relative to C2** | **-15.4** | **5.8** | **-2.6** | **11.0** | **0.02** |

**Table S3C.** – Same as Table S3A but only frontal electrodes.

| Dataset | Coordinate | Name | Estimate | SE | tStat | DF | pValue |
| --- | --- | --- | --- | --- | --- | --- | --- |
| **Dataset 2** | **y** | **C1 - reference** | **21.1** | **4.3** | **4.9** | **5.0** | **0.0045** |
| Dataset 2 | y | C2 - relative to C1 | -1.0 | 7.4 | -0.1 | 8.8 | 0.9 |
| Dataset 2 | y | C3 - relative to C1 | -10.5 | 10.4 | -1.0 | 5.1 | 0.4 |
| **Dataset 2** | **y** | **C2 - reference** | **20.1** | **5.8** | **3.4** | **7.1** | **0.011** |
| Dataset 2 | y | C3 - relative to C2 | -9.4 | 8.9 | -1.1 | 18.5 | 0.3 |
| Dataset 2 | y | C1 - relative to C2 | 1.0 | 7.4 | 0.1 | 8.8 | 0.9 |
| Dataset 2 | z | C1 - reference | 11.7 | 6.6 | 1.8 | 7.5 | 0.11 |
| Dataset 2 | z | C2 - relative to C1 | 6.2 | 6.2 | 1.0 | 6.5 | 0.4 |
| Dataset 2 | z | C3 - relative to C1 | 6.1 | 9.4 | 0.6 | 5.2 | 0.5 |
| **Dataset 2** | **z** | **C2 - reference** | **18.0** | **5.5** | **3.3** | **6.7** | **0.015** |
| Dataset 2 | z | C3 - relative to C2 | -0.2 | 8.2 | 0.0 | 7.8 | 1.0 |
| Dataset 2 | z | C1 - relative to C2 | -6.2 | 6.2 | -1.0 | 6.5 | 0.4 |

**Table S3D.** – Same as Table S3A but only temporal electrodes.

| Dataset | Coordinate | Name | Estimate | SE | tStat | DF | pValue |
| --- | --- | --- | --- | --- | --- | --- | --- |
| **Dataset 2** | **y** | **C1 - reference** | **-24.1** | **5.4** | **-4.4** | **6.2** | **0.0040** |
| Dataset 2 | y | C2 - relative to C1 | -12.8 | 7.0 | -1.8 | 8.1 | 0.10 |
| Dataset 2 | y | C3 - relative to C1 | -4.5 | 9.8 | -0.5 | 7.6 | 0.66 |
| **Dataset 2** | **y** | **C2 - reference** | **-36.9** | **4.0** | **-9.3** | **39.1** | **2E-11** |
| Dataset 2 | y | C3 - relative to C2 | 8.3 | 8.6 | 1.0 | 52.8 | 0.34 |
| Dataset 2 | y | C1 - relative to C2 | 12.8 | 7.0 | 1.8 | 8.1 | 0.10 |
| Dataset 2 | z | C1 - reference | -4.0 | 4.9 | -0.8 | 5.5 | 0.45 |
| Dataset 2 | z | C2 - relative to C1 | 8.6 | 5.4 | 1.6 | 46.8 | 0.11 |
| Dataset 2 | z | C3 - relative to C1 | 12.5 | 10.2 | 1.2 | 6.0 | 0.27 |
| Dataset 2 | z | C2 - reference | 4.6 | 4.7 | 1.0 | 7.4 | 0.36 |
| Dataset 2 | z | C3 - relative to C2 | 3.9 | 10.6 | 0.4 | 5.6 | 0.72 |
| Dataset 2 | z | C1 - relative to C2 | -8.6 | 5.4 | -1.6 | 46.8 | 0.11 |

**Table S4A.** – LME results comparing coordinates of the 3 clusters, Dataset 2, Right hemisphere. Similar to Table S3A but for right-hemisphere electrodes. The significant comparisons were of the y-coordinates of Cluster 2 vs. 1 (p<0.05). See **Figure S5**.

| Dataset | Coordinate | Name | Estimate | SE | tStat | DF | pValue |
| --- | --- | --- | --- | --- | --- | --- | --- |
| Dataset 2 | y | C1 - reference | 2.49 | 7.18 | 0.35 | 7.70 | 0.74 |
| **Dataset 2** | **y** | **C2 - relative to C1** | **-17.17** | **7.02** | **-2.45** | **11.13** | **0.03** |
| Dataset 2 | y | C3 - relative to C1 | -7.00 | 7.52 | -0.93 | 8.64 | 0.38 |
| **Dataset 2** | **y** | **C2 - reference** | **-14.68** | **4.39** | **-3.35** | **3.20** | **0.04** |
| Dataset 2 | y | C3 - relative to C2 | 10.18 | 7.84 | 1.30 | 2.29 | 0.31 |
| **Dataset 2** | **y** | **C1 - relative to C2** | **17.17** | **7.02** | **2.45** | **11.13** | **0.03** |
| Dataset 2 | z | C1 - reference | -8.56 | 5.94 | -1.44 | 6.44 | 0.20 |
| Dataset 2 | z | C2 - relative to C1 | 5.57 | 4.97 | 1.12 | 2.68 | 0.35 |
| Dataset 2 | z | C3 - relative to C1 | 14.76 | 5.99 | 2.46 | 4.09 | 0.07 |
| Dataset 2 | z | C2 - reference | -2.98 | 6.38 | -0.47 | 2.47 | 0.68 |
| Dataset 2 | z | C3 - relative to C2 | 9.18 | 4.69 | 1.96 | 0.74 | 0.36 |
| Dataset 2 | z | C1 - relative to C2 | -5.57 | 4.97 | -1.12 | 2.68 | 0.35 |

**Table S4B.** – Same as Table S4A but electrodes are weighted by reliability. The significant comparisons were of the y-coordinates of Cluster 2 vs. 1, and the z-coordinate of Cluster 3 relative to 1 (ps<0.05).

| Dataset | Coordinate | Name | Estimate | SE | tStat | DF | pValue |
| --- | --- | --- | --- | --- | --- | --- | --- |
| Dataset 2 | y | C1 - reference | 3.7 | 9.3 | 0.4 | 8.1 | 0.70 |
| **Dataset 2** | **y** | **C2 - relative to C1** | **-17.2** | **8.5** | **-2.0** | **72.4** | **0.047** |
| Dataset 2 | y | C3 - relative to C1 | -4.5 | 9.3 | -0.5 | 10.9 | 0.64 |
| Dataset 2 | y | C2 - reference | -13.4 | 7.4 | -1.8 | 1.6 | 0.24 |
| Dataset 2 | y | C3 - relative to C2 | 12.7 | 6.3 | 2.0 | 1.3 | 0.25 |
| **Dataset 2** | **y** | **C1 - relative to C2** | **17.2** | **8.5** | **2.0** | **72.4** | **0.047** |
| Dataset 2 | z | C1 - reference | -8.0 | 6.9 | -1.2 | 6.5 | 0.29 |
| Dataset 2 | z | C2 - relative to C1 | 6.9 | 7.5 | 0.9 | 2.1 | 0.45 |
| **Dataset 2** | **z** | **C3 - relative to C1** | **16.4** | **6.0** | **2.8** | **7.2** | **0.027** |
| Dataset 2 | z | C2 - reference | -1.1 | 8.1 | -0.1 | 2.6 | 0.90 |
| Dataset 2 | z | C3 - relative to C2 | 9.5 | 5.2 | 1.8 | 0.4 | 0.54 |
| Dataset 2 | z | C1 - relative to C2 | -6.9 | 7.5 | -0.9 | 2.1 | 0.45 |

**Table S4C.**
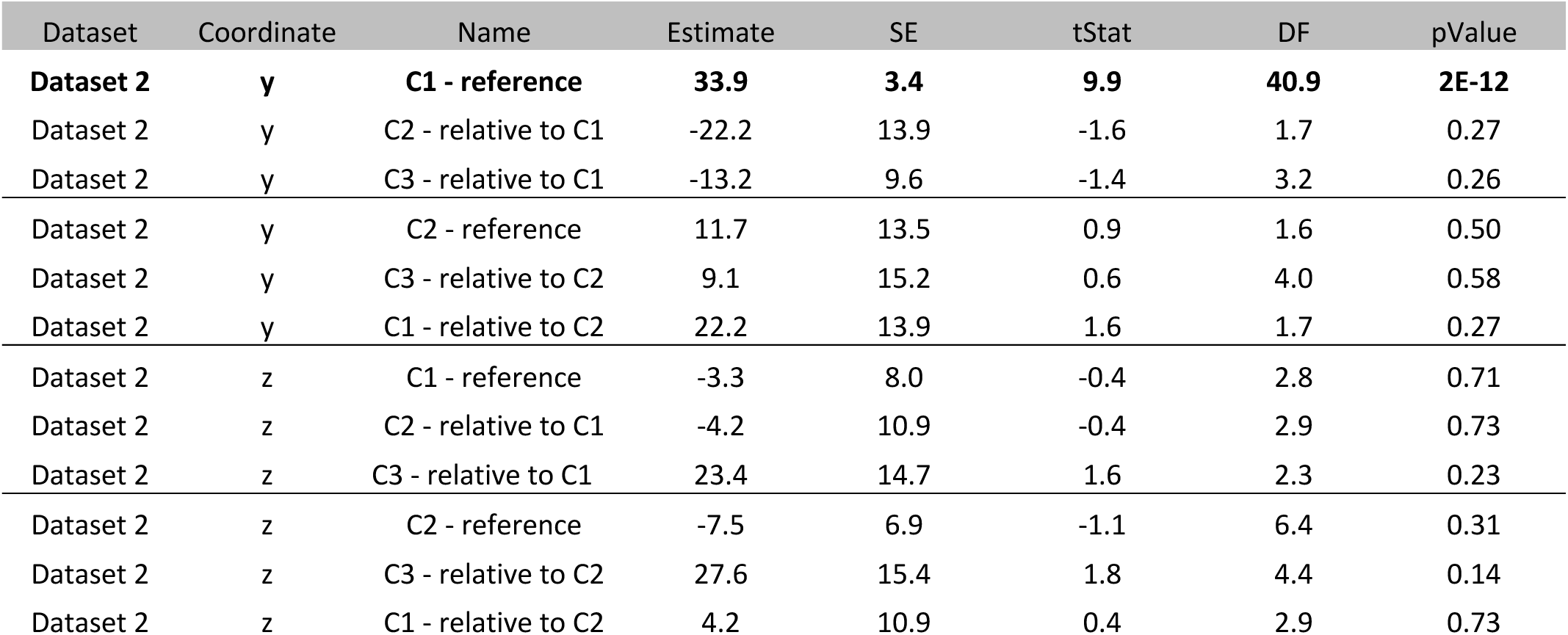
– Same as Table S4A but only frontal electrodes. No significant comparisons.

| Dataset | Coordinate | Name | Estimate | SE | tStat | DF | pValue |
| --- | --- | --- | --- | --- | --- | --- | --- |
| **Dataset 2** | **y** | **C1 - reference** | **33.9** | **3.4** | **9.9** | **40.9** | **2E-12** |
| Dataset 2 | y | C2 - relative to C1 | -22.2 | 13.9 | -1.6 | 1.7 | 0.27 |
| Dataset 2 | y | C3 - relative to C1 | -13.2 | 9.6 | -1.4 | 3.2 | 0.26 |
| Dataset 2 | y | C2 - reference | 11.7 | 13.5 | 0.9 | 1.6 | 0.50 |
| Dataset 2 | y | C3 - relative to C2 | 9.1 | 15.2 | 0.6 | 4.0 | 0.58 |
| Dataset 2 | y | C1 - relative to C2 | 22.2 | 13.9 | 1.6 | 1.7 | 0.27 |
| Dataset 2 | z | C1 - reference | -3.3 | 8.0 | -0.4 | 2.8 | 0.71 |
| Dataset 2 | z | C2 - relative to C1 | -4.2 | 10.9 | -0.4 | 2.9 | 0.73 |
| Dataset 2 | z | C3 - relative to C1 | 23.4 | 14.7 | 1.6 | 2.3 | 0.23 |
| Dataset 2 | z | C2 - reference | -7.5 | 6.9 | -1.1 | 6.4 | 0.31 |
| Dataset 2 | z | C3 - relative to C2 | 27.6 | 15.4 | 1.8 | 4.4 | 0.14 |
| Dataset 2 | z | C1 - relative to C2 | 4.2 | 10.9 | 0.4 | 2.9 | 0.73 |

**Table S4D.** – Same as Table S4A but only temporal electrodes. No significant comparisons.

| Dataset | Coordinate | Name | Estimate | SE | tStat | DF | pValue |
| --- | --- | --- | --- | --- | --- | --- | --- |
| **Dataset 2** | **y** | **C1 - reference** | **-17.5** | **5.6** | **-3.1** | **5.8** | **0.021** |
| Dataset 2 | y | C2 - relative to C1 | -3.7 | 7.5 | -0.5 | 13.4 | 0.63 |
| Dataset 2 | y | C3 - relative to C1 | -6.4 | 6.7 | -1.0 | 28.7 | 0.35 |
| **Dataset 2** | **y** | **C2 - reference** | **-21.2** | **6.4** | **-3.3** | **2.9** | **0.048** |
| Dataset 2 | y | C3 - relative to C2 | -2.7 | 6.1 | -0.4 | 6.6 | 0.68 |
| Dataset 2 | y | C1 - relative to C2 | 3.7 | 7.5 | 0.5 | 13.4 | 0.63 |
| Dataset 2 | z | C1 - reference | -10.7 | 7.7 | -1.4 | 3.3 | 0.25 |
| Dataset 2 | z | C2 - relative to C1 | -4.1 | 9.4 | -0.4 | 3.0 | 0.69 |
| Dataset 2 | z | C3 - relative to C1 | -1.6 | 8.9 | -0.2 | 5.4 | 0.87 |
| Dataset 2 | z | C2 - reference | -14.8 | 4.9 | -3.0 | 0.6 | 0.30 |
| Dataset 2 | z | C3 - relative to C2 | 2.5 | 5.9 | 0.4 | 3.1 | 0.70 |
| Dataset 2 | z | C1 - relative to C2 | 4.1 | 9.4 | 0.4 | 3.0 | 0.69 |

**Table S5.** – LME results comparing temporal receptive windows (TRW) of the 3 clusters, Dataset 1. All estimates from the linear mixed-effects model (LME) regressing the estimated temporal receptive window (TRW) size (<u>Methods</u>) on the categorical variable of cluster (3 levels) grouped by the random variable of participant. Model formula: *trw ∼ cluster + (cluster|participant).* The Satterthwaite Method was used to estimate the degrees of freedom (DF) due to our small sample size. Details are similar to Table S1A. All comparisons were statistically significant: Cluster 2 had a smaller TRW compared to Cluster 1, and Cluster 3 had the smallest trw compared to both other clusters (all ps<0.01). See **Figure 4**.

| Name | Estimate | SE | tStat | DF | pValue |
| --- | --- | --- | --- | --- | --- |
| **C1 - reference** | **6.5** | **0.5** | **12.6** | **5.1** | **5.2E-05** |
| **C2 - relative to C1** | **-2.5** | **0.6** | **-4.0** | **3.6** | **0.020** |
| **C3 - relative to C1** | **-5.1** | **0.6** | **-8.1** | **8.7** | **2.6E-05** |
| **C2 - reference** | **4.0** | **0.5** | **7.9** | **4.3** | **0.0010** |
| **C3 - relative to C2** | **-2.5** | **0.6** | **-4.0** | **8.0** | **0.0040** |
| **C1 - relative to C2** | **2.5** | **0.6** | **4.0** | **3.6** | **0.020** |

**Table S6:** LME results comparing temporal receptive windows (TRW) of the 3 clusters, Dataset 1, due to stimulus presentation rate. Similar to Table S5 but here we add the categorical variable Rate, representing stimulus presentation rate, with two levels: fast (450 ms inter-stimulus interval, n=3) or slow (700 ms inter-stimulus interval, n=3). Model formula: trw ∼ Cluster*Rate + (Cluster*Rate|participant). The model was coded such that one level from each categorical variable was coded as the reference (intercept, whose estimate was compared to 0 for statistical testing). All other levels of the Cluster variable were modeled relative to the reference, and other levels of Rate were modeled relative to the corresponding estimate (see variable names in table). We ran 4 models (LME 1-4) that differed in the order of the levels of the categorical variables, such that at each model a different level was coded as the reference. This allowed us to statistically compare all possible pairs of categories, using the LME stats output (Columns 4-6). DF were estimated using the Satterthwaite approximation. Overall, all models show a negative trend of TRW by Cluster for both presentation rates (smaller TRWs for C3 relative to C2 and for C2 relative to C1). Rate affected only the TRW of Cluster 1 (larger TRW for C1 with slow relative to fast presentation rates) but not of Clusters 2 and 3. The overall main effects of the interaction between Cluster and Rate are not significant due to an additional ANOVA (**Table S7**).

| Name | Estimate | SE | tStat | DF | pValue |
| --- | --- | --- | --- | --- | --- |
| **C1, fast - reference** | **2.61** | **0.31** | **8.43** | **2.17** | **0.011** |
| **C2, fast - relative to ref** | **-1.02** | **0.27** | **-3.79** | **3.36** | **0.026** |
| C3, fast - relative to ref | -2.11 | 0.50 | -4.20 | 1.75 | 0.065 |
| **C1, slow - relative to ref** | **2.64** | **0.48** | **5.49** | **4.54** | **0.004** |
| C2, slow - relative to C2, fast | -1.30 | 0.98 | -1.33 | 2.38 | 0.296 |
| C3, slow - relative to C3, fast | -1.64 | 0.88 | -1.87 | 3.76 | 0.139 |
| **C1, slow - reference** | **5.25** | **0.37** | **14.35** | **2.43** | **0.002** |
| C2, slow - relative to ref | -2.32 | 0.94 | -2.47 | 2.04 | 0.130 |
| **C3, slow - relative to ref** | **-3.75** | **0.72** | **-5.21** | **2.22** | **0.028** |
| **C1, fast - relative to C1, slow** | **-2.64** | **0.48** | **-5.49** | **4.54** | **0.004** |
| C2, fast - relative to C2, slow | 1.30 | 0.98 | 1.33 | 2.38 | 0.296 |
| C3, fast - relative to C3, slow | 1.64 | 0.88 | 1.87 | 3.76 | 0.139 |
| **C2, fast - reference** | **1.60** | **0.21** | **7.43** | **3.05** | **0.005** |
| C3, fast - relative to ref | -1.09 | 0.44 | -2.50 | 3.82 | 0.069 |
| **C1, fast - relative to ref** | **1.02** | **0.27** | **3.79** | **3.36** | **0.026** |
| C2, slow - relative to ref | 1.34 | 0.68 | 1.97 | 2.46 | 0.163 |
| C3, slow - relative to C3, fast | -0.34 | 0.82 | -0.42 | 2.58 | 0.706 |
| C1, slow - relative to C1, fast | 1.30 | 0.98 | 1.33 | 2.38 | 0.296 |
| C3, fast - reference | 0.51 | 0.37 | 1.39 | 29.71 | 0.176 |
| C1, fast - relative to ref | 2.11 | 0.50 | 4.20 | 1.75 | 0.065 |
| C2, fast - relative to ref | 1.09 | 0.44 | 2.50 | 3.82 | 0.069 |
| C3, slow - relative to ref | 0.99 | 0.67 | 1.48 | 4.46 | 0.207 |
| C1, slow - relative to C1, fast | 1.64 | 0.88 | 1.87 | 3.76 | 0.139 |
| C2, slow - relative to C2, fast | 0.34 | 0.82 | 0.42 | 2.58 | 0.706 |

**Table S7.** ANOVA for the LME results presented in Table S6. ANOVA for LME was run on all 4 LME models presented in **Table S5**. NaN as a p-value indicated that there were not sufficient degrees of freedom (DF) to evaluate the statistical effect. Importantly, the interaction between Cluster and Rate did not reach significance.

| Name | FStat | DF1 | DF2 | pValue |
| --- | --- | --- | --- | --- |
| **Intercept (C1, fast)** | **71.1** | **1** | **2.2** | **0.01** |
| Cluster | 10.8 | 2 | 0 | NaN |
| **Rate (C1, slow)** | **30.2** | **1** | **4.5** | **0.004** |
| Cluster:Rate | 1.8 | 2 | 2.7 | 0.3 |
| **Intercept (C1, slow)** | **205.8** | **1** | **2.4** | **0.002** |
| Cluster | 14.7 | 2 | 0 | NaN |
| **Rate (C1, fast)** | **30.2** | **1** | **4.5** | **0.004** |
| Cluster:Rate | 1.8 | 2 | 2.7 | 0.321 |
| **Intercept (C2, fast)** | **55.2** | **1** | **3.1** | **0.005** |
| **Cluster** | **10.8** | **2** | **3.5** | **0.032** |
| Rate (C2, slow) | 3.9 | 1 | 2.5 | 0.163 |
| Cluster:Rate | 1.8 | 2 | 2.5 | 0.329 |
| Intercept (C3, fast) | 1.9 | 1 | 29.7 | 0.176 |
| Cluster | 10.8 | 2 | 2.1 | 0.079 |
| Rate (C3, slow) | 2.2 | 1 | 4.5 | 0.207 |
| Cluster:Rate | 1.8 | 2 | 2.6 | 0.326 |

**Table S8.**
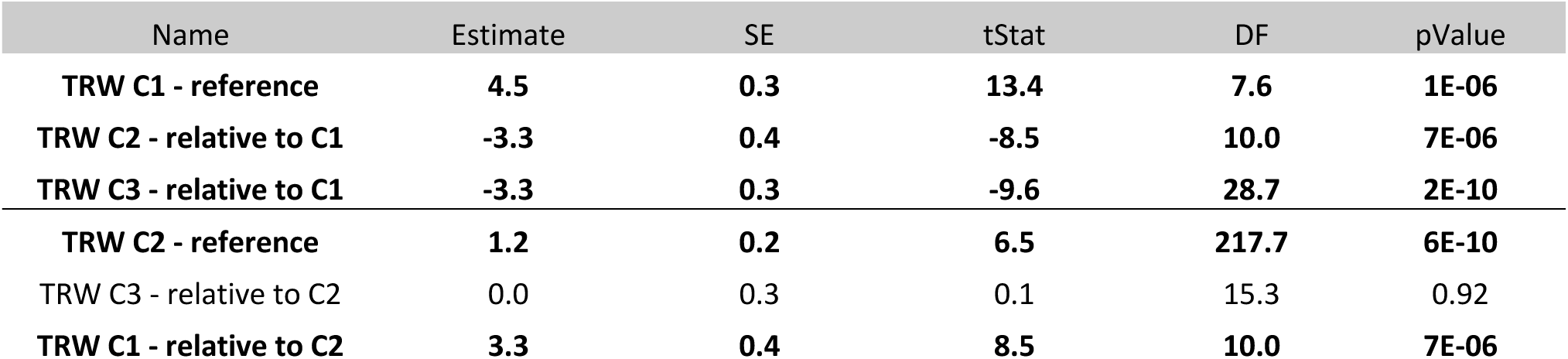
– LME results comparing temporal receptive windows (TRW) of the 3 clusters, Dataset 2. Similar to Table S5, but for Dataset 2 using the first 8 words per each trial. The TRW of C2 is smaller than C1 (p<0.0001) but the same as of C3. See **Figure S10A-B**.

| Name | Estimate | SE | tStat | DF | pValue |
| --- | --- | --- | --- | --- | --- |
| **TRW C1 - reference** | **4.5** | **0.3** | **13.4** | **7.6** | **1E-06** |
| **TRW C2 - relative to C1** | **-3.3** | **0.4** | **-8.5** | **10.0** | **7E-06** |
| **TRW C3 - relative to C1** | **-3.3** | **0.3** | **-9.6** | **28.7** | **2E-10** |
| **TRW C2 - reference** | **1.2** | **0.2** | **6.5** | **217.7** | **6E-10** |
| TRW C3 - relative to C2 | 0.0 | 0.3 | 0.1 | 15.3 | 0.92 |
| **TRW C1 - relative to C2** | **3.3** | **0.4** | **8.5** | **10.0** | **7E-06** |

**Table S9.** – LME results comparing temporal receptive windows (TRW) of the 3 clusters, Dataset 2, using 8 words, electrodes assigned to clusters by similarity to Dataset 1 cluster centers. Similar to Table S8, but here the grouping of electrodes to the 3 clusters was done by assigning each electrode in Dataset 2 to a cluster by its highest correlation with the average cluster response profiles from Dataset 1. All comparisons were statistically significant: Cluster 2 had a smaller TRW compared to Cluster 1, and Cluster 3 had the smallest TRW compared to both other clusters (all ps<0.001). See **Figure S10C-D**.

| Name | Estimate | SE | tStat | DF | pValue |
| --- | --- | --- | --- | --- | --- |
| **TRW C1 – reference** | **4.50** | **0.29** | **15.5** | **10.2** | **2.0E-08** |
| **TRW C2 – relative to C1** | **-2.38** | **0.29** | **-8.2** | **8.1** | **3.3E-05** |
| **TRW C3 – relative to C1** | **-3.49** | **0.32** | **-10.8** | **14.1** | **3.4E-08** |
| **TRW C2 – reference** | **2.12** | **0.27** | **7.8** | **11.2** | **7.2E-06** |
| **TRW C3 – relative to C2** | **-1.11** | **0.31** | **-3.6** | **15.6** | **2.5E-03** |
| **TRW C1 – relative to C2** | **2.38** | **0.29** | **8.2** | **8.1** | **3.3E-05** |

